# Regulation of essential hepatocyte functions and identity by super-enhancers in health and disease

**DOI:** 10.1101/2025.06.04.657826

**Authors:** Tao Lin, Chenjun Huang, Chenhao Tong, Wentao Xu, Han Wang, Shanshan Wu, Wenzhe Zhang, Yujia Li, Hui Liu, Chen Shao, Rui Liu, ShansHan Wang, Stefan Munker, Hanno Niess, Christoph Meyer, Roman Liebe, Matthias P Ebert, Chunfang Gao, Steven Dooley, Huiguo Ding, Li Zuo, Hua Wang, Hong-Lei Weng

## Abstract

Physiologically, hepatocytes perform an estimated 500 essential liver functions to ensure systemic homeostasis. The mechanisms how hepatocytes maintain these numerous crucial functions in response to physiological and pathophysiological challenges are largely unknown. In this study, we found that vital liver function genes (e.g., *HNF4A*, *ALB*, coagulation factors, and *SLC2A2*) in hepatocytes possess super-enhancers. Interfering with the super-enhancers by dCas9-sgRNA remarkably decreases the expression of these genes. Quiescent liver progenitor cells (LPCs) do not have active super-enhancers in vital liver function genes. In acute liver failure (ALF) caused by massive hepatic necrosis, activated LPCs perform liver function and differentiate into hepatocytes to rescue patients’ lives. We show that activated LPCs gradually form super-enhancers in liver function genes (e.g., *HNF4A*, *ALB*, and coagulation factors) during proliferation and differentiation. These super-enhancers are regulated by FOXA2 through the recruitment of HNF4 a, p300 and Mediators in most liver function genes, such as *ALB* and coagulation factors. However, super-enhancers in some genes (e.g., *SLC2A2*) are FOXA2-indenpendent. Hepatic FOXA2 expression is closely associated with the clinical outcome of ALF. In a cohort of ALF patients, robust FOXA2 is observed only in the surviving patients. Collectively, super-enhancers represent a critical epigenetic regulatory mechanism that enables rapidly enhanced transcription of liver function genes in response to pathophysiological challenges. In urgent clinical syndrome such as ALF, LPCs rescue liver function by rapid formation of super-enhancers in vital genes.

---

The liver is the largest metabolic organ in the human body and performs an estimated total of around 500 functions: Carbohydrate, lipid, and amino acid metabolism, the synthesis of essential proteins such as *ALB* and coagulation factors; and the production of other biochemicals necessary for digestion and growth (e.g., bile and growth factors), as well as detoxification of the organism ^1, 2 3^. All these functions are carried out by hepatocytes and are crucial for human systemic homeostasis ^3^. How hepatocytes perform such a large number and wide variety of functions in response to physiological needs and pathophysiological challenges is still an exciting and relevant question.

In the past decade, the role of super-enhancers in controlling cell state and identity has been uncovered ^4^. Super-enhancers are putative enhancer clusters in the vicinity of genes, showing unusually high levels of enhancer activity. They are occupied by master transcription factors, co-factors, chromatin regulators and core transcription machinery, including master transcription factors such as Oct4, Sox2 and Nanog; the Mediators; co-activators (also known as pioneer factors); insulators like CTCF and cohesin; chromatin regulators such as Smc1 and Brg1; and the histone acetyltransferases like CBP/p300 ^4^. The representative structure of super-enhancers has been clarified in embryonic stem cells (ESCs). Master transcription factors recruit Mediators and cohesin to organize loop structures between promoters and enhancers. Within these loops, master transcription factors open chromatin and render enhancers accessible, which is characterized by enriched H3K27ac. In collaboration with chromatin regulators and histone acetyltransferases, master transcription factors assemble enhancer clusters, which are bound by multiple transcription activators to regulate specific genes ^4^. To date, the structure and function of super-enhancers have been well characterized ESCs and cancer cells ^4^.

In the liver, few studies have investigated the effects of super-enhancers on the expression of vital liver function genes in hepatocytes. To date, an LRH1-driven super-enhancer-associated network is reported to regulate hepatocyte identity ^5^. A few recent studies focus on the role of super-enhancers in regulating liver cancer-associated signaling pathways and oncogenes ^6^. Given that super-enhancers possess unusually high levels of enhancer activity, we speculate whether these epigenetic structures play a critical role in regulating vital liver function genes in response to different physiological changes. As the largest metabolic organ in the human body, the liver is permanently exposed to metabolites and toxins. Therefore, hepatocytes must have evolved different regulatory mechanisms that ensure the transcription of vital liver function genes as needed.

Beyond physiological conditions, the liver may suffer from massive hepatic necrosis (MHN) and progresses into acute liver failure (ALF), an urgent clinical syndrome with high mortality, under an extremely insults ^7–9^. In this situation, the severely damaged and functionally compromised liver relies on rapidly activated liver progenitor cells (LPCs) to perform essential liver functions, to maintain systemic homeostasis and thus ensure patients’ survival ^7, 10^. LPCs are the smallest cholangiocytes localized in the Canals of Hering and the smallest branches of the biliary tree ^11–13^. Morphologically, LPCs are cells measuring less than 6 μm, significantly smaller than hepatocytes, which range from 20 to 40 μm ^12^. A key question is how such small LPCs rapidly and adequately perform hepatocyte functions to maintain liver homeostasis in ALF patients. Would super-enhancers control the expression of liver function genes in LPCs under such emergency conditions?

In this study, we confirm the critical role of super-enhancers in the transcription of the liver function genes in hepatocytes. Vital liver function genes (e.g., *HNF4A*, *ALB*, coagulation factors, and *SLC2A2*) in hepatocytes possess super-enhancers. Interfering with the activity of super-enhancers by CRISPR-dCas9-sgRNA remarkably inhibits the expression of these essential hepatocyte identity genes. In contrast to hepatocytes, quiescent LPCs do not form super-enhancers in vital liver function genes under normal circumstances. However, activated LPCs form super-enhancers in essential liver function genes in emergency situations to support the differentiation toward hepatocytes. Pioneer factor FOXA2 is a key regulator that controls super-enhancer activation in most liver function genes (e.g., *ALB* and coagulation factors) by recruiting HNF4 α, histone acetyltransferase p300, and Mediators to super-enhancers and maintaining super-enhancer loops. However, super-enhancers in other genes (e.g., *SLC2A2*) are FOXA2-indenpendent. In MHN-dependent ALF, hepatic FOXA2 expression is closely associated with the clinical outcome of the disease. In a cohort of ALF patients, robust FOXA2 expression was observed only in surviving patients.

## Results

### Liver function genes in hepatocytes possess super-enhancers

To identify super-enhancer in the liver, we analyzed ChIP-seq databases from human livers, HepG2 cells, and mouse livers. Super-enhancer was defined using the standard ROSE algorithms based on H3K27ac signals ^14^. Based on enrichment of H3K27ac signals, 1433, 1680, 1079, 1427 and 880 super-enhancers were identified in two human livers, HepG2 cells, and two mouse livers, respectively (**Figure 1A** and **Figure S1A**). A large portion of liver function genes possess super-enhancers, including master hepatic transcriptional factors (*HNF4A*, *CEBPA* and *HNF1B*), coagulation factors (*F5*, *F7*, *F10*, *F11*, and *F12*), *ALB*, cytochrome P450 family member (*CYP2E1* and *CYP8B1*), alcohol dehydrogenase *ADH4* and Aldehyde dehydrogenase *ALDH2*, Apolipoprotein B *APOB*, and the glucose transporter gene *SLC2A2* (**Figure 1A** and **Figure S1A**). KEGG analyses revealed that these genes were enriched in critical liver functional categories, including coagulation cascade, biosynthesis, drug/energy metabolism (**Figure S1B**). **Table S1** summarizes transcription factors and liver function genes with super-enhancers in either human liver and HepG2 cells. Compared to typical enhancers, these super-enhancers exhibited significantly elevated H3K27ac signals in livers and HepG2 cells (**Figure 1B** and **Figure S1C**).

**Figure 1.**
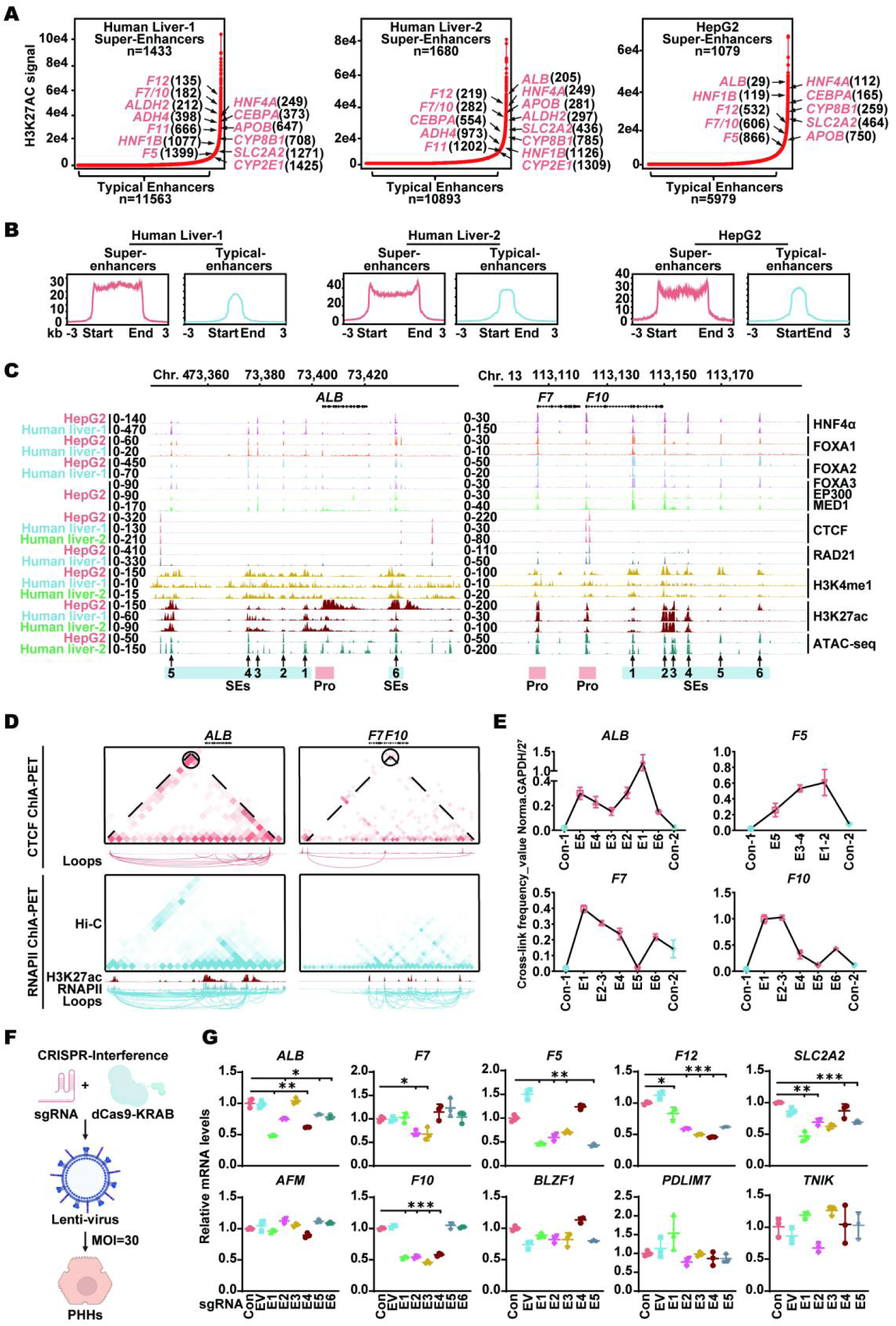
Identification and functional validation of liver function gene-associated super-enhancers in hepatocytes. (A) Super-enhancers (SEs) were identified based on normalized H3K27ac ChIP-seq signal intensity using the ROSE algorithm in independent two human livers and HepG2 cells. Super enhancers are defined as the enhancers above the inflection point of the enhancer rank curve, while the rest are classified as typical enhancers (TEs). Key liver genes associated with SE are marked in red with their enhancer rank. (B) Average H3K27ac ChIP-seq signal is centered around SEs and TEs in human livers and HepG2 cells. The signal is plotted from 3 kb upstream to 3 kb downstream of the enhancer regions. (C) Genomic browser tracks show the occupancy of indicated proteins and chromatin accessibility (ATAC-seq) at representative super-enhancers associated loci (*ALB*, *F7*, and *F10*) in HepG2 cells (red) and human liver tissues (blue and green). Arrows denote identified SE regions, and shaded boxes highlight regions of super-enhancers and promoters. (D) Hi-C, ChIA-PET interactions, and ChIP-seq maps of the *ALB*, *F7*, and *F10* loci in HepG2 cells. Top (CTCF ChIA-PET): CTCF-mediated chromatin loops are shown in red. Hi-C contact maps at 10 kb (ALB) or 5 kb (*F7* and *F10*) resolution reveal local topologically associated domains (TADs), while CTCF ChIA-PET loops highlight the structural organization and long-range interactions within these regions. Bottom (RNAPII ChIA-PET): RNAPII-mediated enhancer-promoter loops are shown in blue. H3K27ac ChIP-seq tracks (red) indicate active enhancer regions. RNAPII ChIA-PET maps reveal regulatory loops connecting super-enhancers and target gene promoters, supporting transcriptional activation. Dashed triangles outline TADs based on Hi-C data. Rings below each contact map represent high-confidence interactions detected by ChIA-PET. (E) 3C-PCR assays validate interactions between candidate enhancer and promoter regions of the indicated genes in primary human hepatocytes. (F) A schematic of the CRISPR interference (CRISPRi) strategy targeting super-enhancers via dCas9-KRAB lentiviral delivery in primary human hepatocytes is presented. (G) Expression levels of mRNA for the indicated genes following control sgRNA (Con), empty vector (EV), or CRISPRi-mediated super-enhancers repression in primary human hepatocytes are shown. Peptidylprolyl isomerase A (PPIA) expression, measured by qRT-PCR, was used as a normalization control. Each point represents an independent biological replicate. Data are presented as mean ± standard error of the mean (SEM) from three independent experiments. P values were calculated using a two-sided Student’s t-test; *P < 0.05, **P < 0.01, ***P < 0.001.

Next, we analyzed super-enhancers associated with the following genes: *ALB*, *SLC2A2*, *CYP8B1*, *HNF4A*, *APOB*, and coagulation factors *F5*, *F7*, *F10*, and *F12*. As shown in **Figure 1C and Figure S2A**, H3K27ac-defined super-enhancers were located upstream of the *ALB* and *APOB* genes, downstream of the *CYP8B1*, *F7*, *F10*, and *F12* genes, and within the gene bodies of *HNF4A*, *F5* and *SLC2A2*. In addition to enriched H3K27ac, these super-enhancers showed occupancy of other enhancer-associated components such as H3K4me1, EP300, and MED1, as well as hepatic master transcription factors (HNF4α, FOXA1, FOXA2, and FOXA3) (**Figure 1C** and **Figure S2A**).

It is well established that DNA regions containing super-enhancers forms chromatin loops to interact with gene promoters, which are confined within cohesin- and CTCF-defined topologically associated domains (TAD) ^15–18^. In HepG2 cells and human livers, ChIP-seq analyses revealed co-bindings of CTCF and RAD21 (a cohesin subunit), at these sites (**Figure 1C and Figure S2A**). ATAC-seq further confirmed accessible chromatin at super-enhancer regions (**Figure 1C and Figure S2A**).

We also performed ChIA-PET to analyze 3D chromatin interaction between the promoter and super-enhancers on the aforementioned liver function genes in HepG2 cells. Integration of CTCF ChIA-PET, Hi-C, and ChIP-seq datasets identified TADs encompassing promoters and super-enhancers in these genes (**Figure 1D** and **Figure S3**). RNAPII ChIA-PET revealed high interaction frequencies within TADs, including frequent promoter-super-enhancer contacts (**Figure 1D** and **Figure S3**). Co-enrichments of RNAPII and H3K27ac at super-enhancer sites further support their role in active transcription (**Figure 1D** and **Figure S3**).

Subsequently, we performed chromosome conformation capture-qPCR (3C-qPCR) to detect interactions between promoters and super-enhancers of *ALB*, *F5*, *F7*, and *F10* in primary human hepatocytes. Except for super-enhancer region E5 of the *F7/F10* locus, interaction frequencies between promoters and super-enhancers within chromatin loops were significantly higher than those outside loops (**Figure 1E**). Collectively, these results validate the presence of super-enhancers associated with liver function genes in hepatocytes.

### In situ CRISPR-interference of super-enhancers inhibits liver function gene expression in hepatocytes

To clarify the effects of super-enhancers on liver function gene expression (e.g., *ALB*, *F5*, *F7*, *F10*, *F12*, and *SLC2A2*), we applied CRISPR-dCas9 interference to target different super-enhancers in fresh isolated human primary hepatocytes (**Figure 1F**). With the exception of E3 in the *ALB* super-enhancer, E4 in the *F5* super-enhancer, E5 and E6 in the *F10* super-enhancer, and E4 in the *SLC2A2* super-enhancer, disruption of any other super-enhancer elements significantly reduced RNA expression of *ALB*, *F5*, *F10*, *F12*, and *SLC2A2* (**Figure 1G**). For *F7*, only the disruption of E2 and E3 affected its expression (**Figure 1G**). The CRISPR interference did not alter mRNA levels of *AFM*, *BLZF1*, *PDLIM7*, and *TNIK*, which are located outside the TADs containing *ALB*, *F5*, *F12*, and *SLC2A2* super-enhancers, respectively (**Figure 1G**), indicating the specificity of super enhancers and the importance of chromatin architecture in regulating gene expression.

These results suggest that super-enhancers associated with liver function genes are essential for their transcription.

### Pioneer factors FOXA2 and FOXA3 are essential for the expression of liver function genes with super-enhancers

As essential components within the promoter-enhancer loops containing super-enhancers, pioneer factors maintain accessible chromatin for the binding of additional transcription factors to super-enhancers ^4^. As described above, pioneer factors FOXA1, FOXA2 and FOXA3 are enriched within super-enhancers of liver function genes (**Figure 1C** and **Figure S2**). To elucidate the role of the FOXA family members in regulating the expression and super-enhancer composition of liver function genes, we investigated how the disruption of FOXA genes affects transcription and epigenetic alterations in the livers and hepatocytes.

According to dataset GSE140423, which were established from mice with individual or combined Foxa genes knockout, GO and KEGG pathway analyses identified multiple altered pathways in the livers of mice with *Foxa* triple knockout. **Figure 2A** highlights the pathways with the highest degree values when *Foxa* triple knockout was induced in mouse livers. Based on GO pathway analysis, these changed pathways were associated with cellular responses to toxic substances, blood coagulation, hemostasis, and metabolic processes related to hormones, cholesterol, steroids, fatty acids, and alcohol. They were also linked to molecular functions such as RNA polymerase II (RNAPII)-specific DNA-binding transcription factor binding, alcohol dehydrogenase (NAD+) activity, heparin binding, heme binding, and lipid transporter activity (**Figure 2A**). KEGG analysis further revealed that the significantly altered pathways were associated with the biosynthesis of unsaturated fatty acids and steroid hormones, bile secretion, drug metabolism via cytochrome P450, retinol metabolism, and cascades of complements and coagulation (**Figure 2A**).

**Figure 2.**
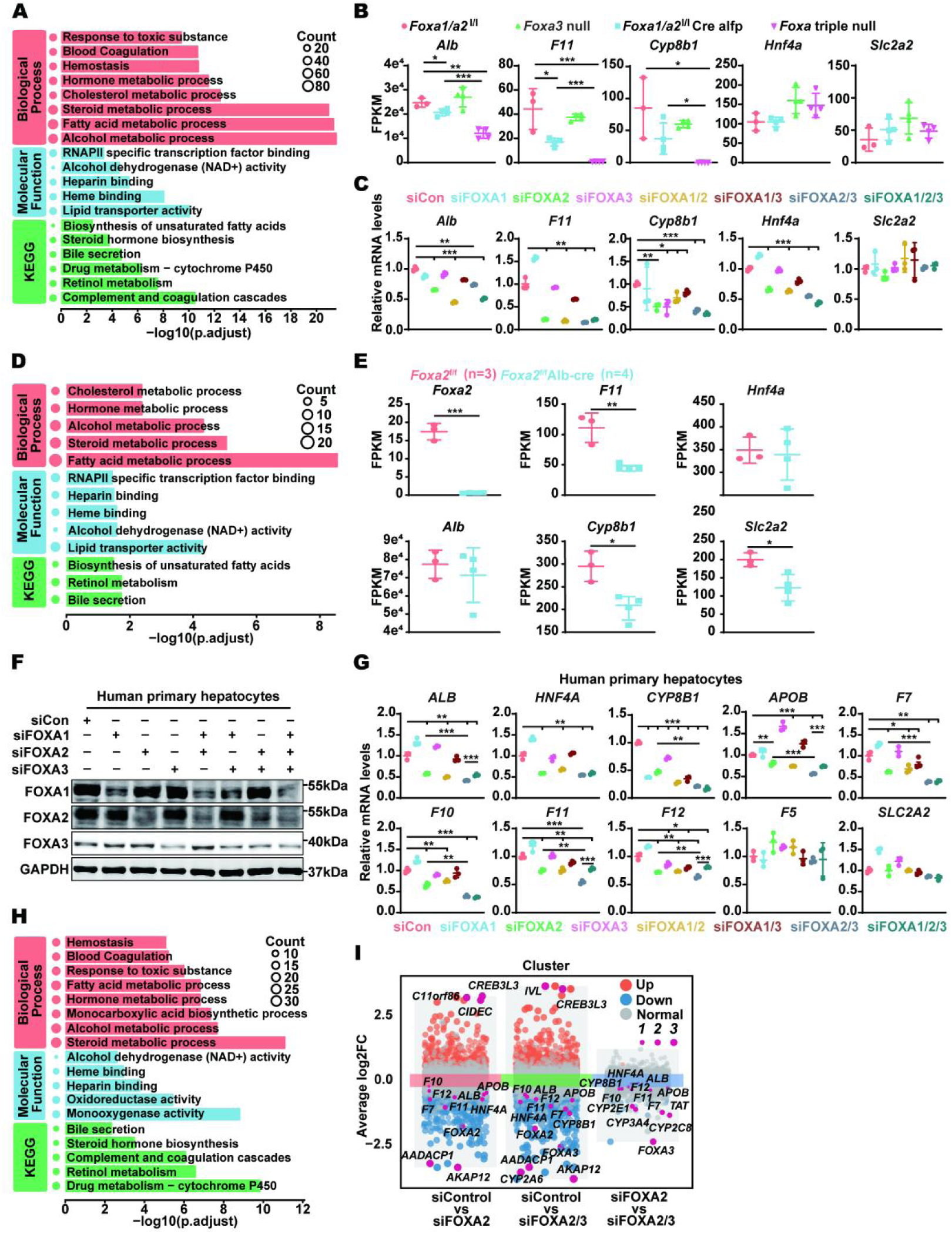
FOXA2 predominantly regulates the expression of most liver function genes with super-enhancers in hepatocytes. (A) A bar plot shows the GO and KEGG term analysis results of the top genes in the livers of *Foxa* triple knockout mice. (B) FPKM was performed to measure *Alb*, *Cyp8b1*, *F11*, *Hnf4a*, and *Slc2a2* transcripts in the livers of mice with different *Foxa* gene disruptions. (C) qPCR was used to analyze liver function gene expression in mouse primary hepatocytes following siRNA-mediated knockdown of individual or combined *Foxa* genes. Data are presented as mean ± standard error of the mean (SEM) from three independent experiments. P values were calculated using a two-sided Student’s t-test; *P < 0.05, **P < 0.01, ***P < 0.001. (D) A bar plot shows the GO and KEGG term analysis results of the top genes for the primary hepatocytes isolated from *Foxa2*^flox/flox^ and *Foxa2*^flox/flox^ Alb-Cre mice. (E) FPKM analysis was performed to measure *Foxa2*, *Alb*, *Cyp8b1*, *F11*, *Hnf4a*, and *Slc2a2* transcripts in mouse primary hepatocytes from *Foxa2*^flox/flox^ and *Foxa2*^flox/flox^ Alb-Cre mice. Data are shown as mean ± SEM. Statistical significance was determined using appropriate statistical tests: *p < 0.05, **p < 0.01, ***p < 0.001. (F) Western blotting shows the expression of FOXA1, FOXA2 and FOXA3 in human primary hepatocytes with different FOXA siRNA. (G) qPCR was used to analyze liver function gene expression in human primary hepatocytes following siRNA-mediated knockdown of individual or combined FOXA genes. Data are presented as mean ± standard error of the mean (SEM) from three independent experiments. P values were calculated using a two-sided Student’s t-test; *P < 0.05, **P < 0.01, ***P < 0.001. (H) A bar plot shows the GO and KEGG term analysis results of the top genes in human primary hepatocytes treated with FOXA2 and/or FOXA3 siRNA. (I) A MA plot displays the differentially expressed genes in human primary hepatocytes after FOXA2 or combined FOXA2/FOXA3 knockdown.

Subsequently, we used FPKM to analyze the transcripts of five genes with super-enhancers (*Alb*, *Cyp8b1*, *F11*, *Hnf4a* and *Slc2a2*) in the livers of mice with different *Foxa* gene disruptions. Compared to *Foxa1*/*a2*^lox/lox^ and *Foxa3* null mice, the transcripts of *Alb*, *Cyp8b1*, and *F11* were significantly reduced in *Foxa1*/*a2*^lox/lox^ Alfp_Cre mice and were further decreased in *Foxa* triple null mice (**Figure 2B**). However, the *Hnf4a* and *Slc2a2* transcripts were not altered in any of the mice with *Foxa* family genes disruption (**Figure 2B**). To further clarify the effects of individual FOXA proteins on liver function gene expression, we used siRNA to silence the expression of individual or combined FOXA in freshly isolated mouse hepatocytes (**Figure 2C**). Knockdown of FOXA2 significantly reduced the mRNA expression of *Alb*, *Cyp8b1*, *Hnf4a* and *F11*, but not *Slc2a2* (**Figure 2C**). Silencing FOXA1 or FOXA3 using RNAi did not reduce the expression of these genes (**Figure 2C**).

Next, we performed RNA sequencing on mouse primary hepatocytes isolated from *Foxa2*^flox/flox^ and *Foxa2*^flox/flox^ Alb-Cre mice. GO analysis showed that, compared to *Foxa2*^flox/flox^ mice, hepatocytes isolated from *Foxa2*^flox/flox^ Alb-Cre mice demonstrated significantly altered pathways associated with responses to cholesterol, hormones, alcohol, steroids, and fatty acid metabolism (**Figure 2D**). Additionally, molecular functions associated with RNAPII-specific DNA-binding transcription factor binding, heparin binding, heme binding, alcohol dehydrogenase (NAD+) activity, and lipid transporter activity were also affected when hepatic *Foxa2* gene was deleted (**Figure 2D**). KEGG analysis further revealed that the significantly altered pathways included the biosynthesis of unsaturated fatty acids, retinol metabolism, and bile secretion (**Figure 2D**). FPKM analysis showed that transcripts of *Cyp8b1*, *F11* and *Slc2a2*, but not *Alb* and *Hnf4a*, were significantly reduced in *Foxa2*^flox/flox^ Alb-Cre mice compared to *Foxa2*^flox/flox^ mice (**Figure 2E**).

In human primary hepatocytes, we analyzed the transcripts of ten genes with super-enhancers (*ALB*, *APOB*, *CYP8B1*, *F5*, *F7*, *F10*, *F11*, *F12*, *HNF4A* and *SLC2A2*) when different FOXA proteins were disrupted. Western blot confirmed that FOXA proteins were effectively knocked down in freshly isolated human primary hepatocytes (**Figure 2F**). Knockdown of FOXA family members, either individually or in combination, did not alter the mRNA expression of *F5* and *SLC2A2* (**Figure 2G**). Knockdown of FOXA1 or FOXA3 using RNAi did not significantly influence the expression of most of the examined genes (**Figure 2G**). FOXA1 siRNA only reduced *CYP8B1* mRNA expression, while silencing FOXA3 slightly decreased the expression of *CYP8B1* and *F12* (**Figure 2G**). In contrast to FOXA1 and FOXA3, silencing FOXA2 significantly reduced the mRNA expression of eight genes: *HNF4A*, *ALB*, *CYP8B1*, *APOB*, *F7*, *F10*, *F11*, and *F12* (**Figure 2G**). Furthermore, simultaneously knocking down both FOXA2 and FOXA3 further reduced the mRNA expression of these genes, except for *HNF4A* (**Figure 2G**). Interestingly, silencing all three FOXA genes simultaneously did not further decrease the expression of these genes (**Figure 2G**). For *ALB*, *APOB*, *F11*, and *F12*, triple knockdown of FOXA genes slightly increased their mRNA expression compared to the double knockdown of FOXA2 and FOXA3 (**Figure 2G**), suggesting a potential antagonistic role of FOXA1 with FOXA2 and FOXA3.

To further clarify the gene profile regulated by FOXA2 and FOXA3 in human hepatocytes, we performed RNA sequencing on human primary hepatocytes treated with or without FOXA2/3 siRNA. GO analysis showed that silencing FOXA2 using RNAi significantly altered pathways associated with homeostasis, blood coagulation, responses to toxic substances, and metabolic processes related to fatty acids, hormones, alcohol, steroids, and monocarboxylic acid biosynthesis (**Figure 2H**). Additionally, molecular functions associated with RNAPII-specific DNA-binding transcription factor binding, heparin binding, heme binding, alcohol dehydrogenase (NAD+) activity, and lipid transporter activity were also affected (**Figure 2H**). KEGG analysis further revealed that the significantly altered pathways included the biosynthesis of unsaturated fatty acids, retinol metabolism, and bile secretion (**Figure 2H**). Subsequently, we used the multi-MA plot-style scatterplot to analyze differential gene expression in PHPs with or without FOXA2 and FOXA3 RNAi. Consistent with the findings based on qPCR analysis (**Figure 2G**), si-FOXA2 reduced the transcripts of genes with super-enhancers, including *ALB*, *APOB*, *HNF4A*, *F7*, *F11*, and *F12*, while double knockdown of FOXA2 and FOXA3 further decreased levels of some transcripts such as *F7*, *F10*, *F11*, and *F12* (**Figure 2I**).

These results suggest that liver function genes with super-enhancers can be classified into three categories: (1) FOXA2-dependent (e.g., *HNF4A*), (2) FOXA2- and FOXA3- dependent (e.g., *ALB* and multiple coagulation factors), and (3) FOXA-independent (e.g., *F5* and *SLC2A2*).

### FOXA2 and FOXA3 are essential for super-enhancers in hepatocytes

Next, we investigated how FOXA2 regulates super-enhancers in hepatocytes. We isolated primary hepatocytes from *Foxa2*^flox/flox^ and *Foxa2*^flox/flox^ Alb-Cre mice and performed H3K27ac ChIP-seq. Based on the enrichment of H3K27ac signals, the number of super-enhancers decreased from 1,332 in normal mouse hepatocytes to 1,154 in *Foxa2*-deficient hepatocytes (**Figure 3A**).

**Figure 3.**
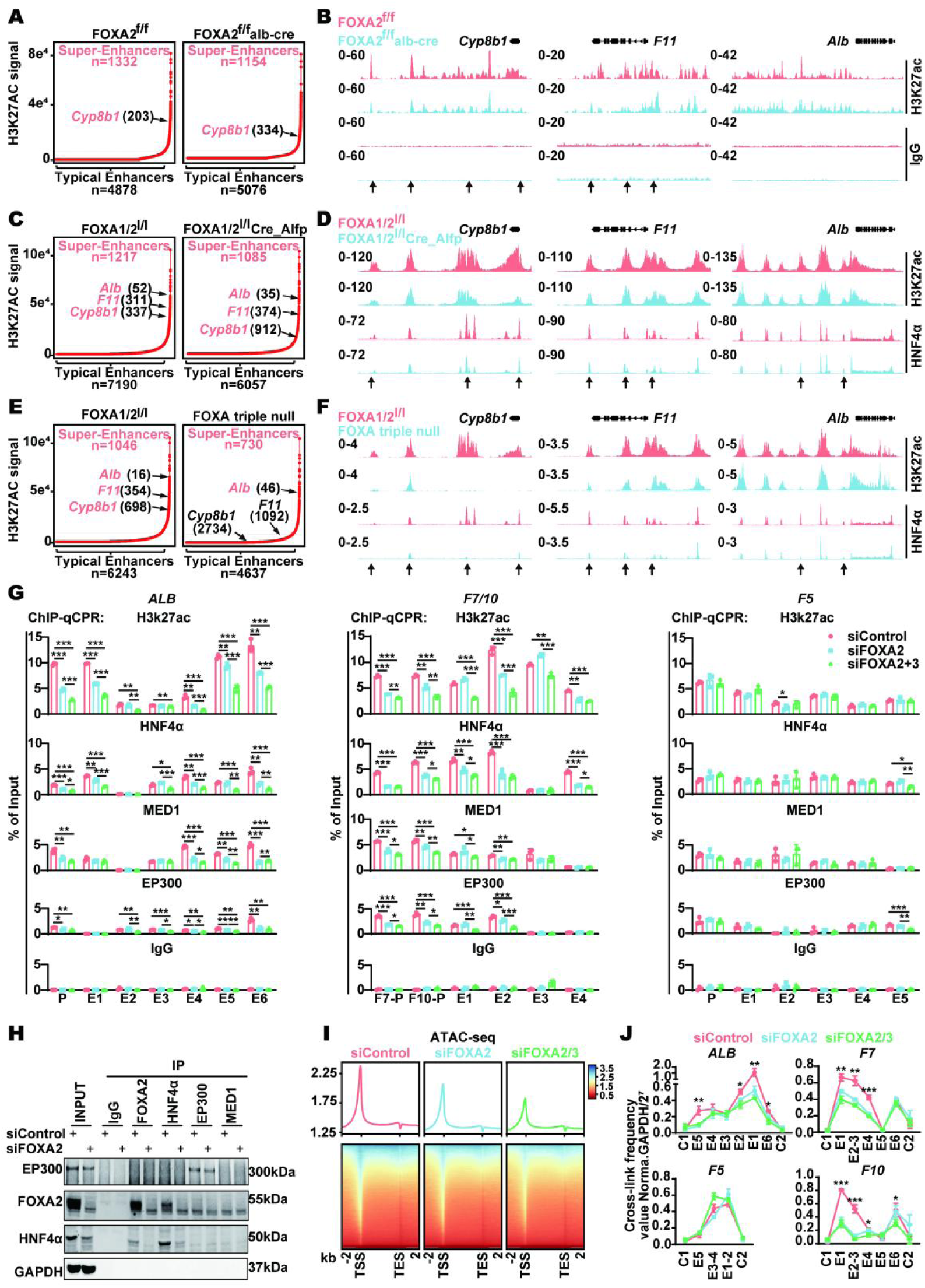
FOXA2 is essential for maintaining super-enhancers and chromatin accessibility in hepatocytes. (A) Super-enhancers were identified based on the normalized H3K27ac ChIP-seq signal using the ROSE algorithm in mouse primary hepatocytes isolated from *Foxa2*^flox/flox^ and *Foxa2*^flox/flox^ Alb-Cre mice. (B) ChIP-seq peaks for H3K27ac and IgG, at the *Alb*, *F11*, and *Cyp8b1* loci were analyzed in primary hepatocytes isolated from *Foxa2*^flox/flox^ (Red) and *Foxa2*^flox/flox^ Alb-Cre (Blue) mice. (**C, E**) Super-enhancers were identified based on the normalized H3K27ac ChIP-seq signal in livers of *Foxa1*/*a2*^lox/lox^, *Foxa1*/*a2*^lox/lox^ Alfp_Cre, and *Foxa* triple null mice (datasets GSE124384 and GSE140423). (**D, F**) ChIP-seq peaks of H3K27ac and HNF4α at the Alb, *F11*, and *CYP8B1* loci were analyzed in livers isolated from *Foxa1*/*a2*^lox/lox^, *Foxa1*/*a2*^lox/lox^ Alfp_Cre, and *Foxa* triple null mice. (G) ChIP-qPCR was performed to examine H3K27ac, HNF4α, EP300, MED1 and IgG occupancy at the promoters (P) and super-enhancers of *ALB*, *F7*, *F10* and *F5* in primary human hepatocytes treated with siRNA-FOXA2 or siRNA-FOXA2/3. Data are shown as mean ± SEM. Statistical significance was determined using appropriate statistical tests: *p < 0.05, **p < 0.01, ***p < 0.001. (H) Co-immunoprecipitation was performed to measure the interaction between FOXA2, HNF4 α, EP300, and MED1 in primary human hepatocytes with or without FOXA2 siRNA treatment. (I) Normalized read distribution profiles of ATAC-seq are presented for human primary hepatocytes treated with siRNA-FOXA2 or siRNA-FOXA2/3. (J) 3C-PCR assays were performed to validate interactions between super-enhancers and promoter regions of the indicated genes in primary human hepatocytes treated with siRNA-FOXA2 or siRNA-FOXA2/3 knockdown. Data are shown as mean ± SEM. Statistical significance was determined using appropriate statistical tests: *p < 0.05, **p < 0.01, ***p < 0.001.

In the absence of *Foxa2*, H3K27ac signal intensity was significantly reduced in the promoters and super-enhancers of the *F11* and the *Cyp8b1* genes but did not affect the *Alb* gene (**Figure 3B**). Although *Cyp8b1* retained its status as a gene with super-enhancers in *Foxa2* deficient hepatocytes, its ranking was lower compared to that in normal hepatocytes (**Figure 3A**). **Table S2** summarizes the genes that lost super-enhancers when *Foxa2* was knocked out.

Subsequently, we analyzed super-enhancers in the livers of *Foxa1*/*a2*^lox/lox^ mice, *Foxa1*/*a2*^lox/lox^ Alfp_Cre mice, and *Foxa* triple null mice using the GSE124384 and GSE140423 database. ChIP-seq for H3K27ac revealed a reduction in the number of super-enhancers in the livers of *Foxa1*/*a2* knockout mice, decreasing from 1,217 to 1,085 (**Figure 3C**). Notably, the super-enhancer rankings of *F11* and *Cyp8b1* were significantly reduced, whereas the ranking of the *Alb* gene showed a slight increase (**Figure 3C**). In addition, the enrichment of H3K27ac in the super-enhancers of *F11*, *Cyp8b1*, and *Alb* genes decreased in *Foxa1*/a*2* knockout mice (**Figure 3D**). Furthermore, HNF4α protein binding to the promoter and/or super-enhancer regions of these genes was substantially reduced in the absence of the *Foxa1* and *Foxa2* genes (**Figure 3D**).

Compared to *Foxa1*/a*2* knockout mice, the number of super-enhancers in *Foxa* triple-knockout mice were reduced from 1,046 to 730 (**Figure 3E**). In the absence of all FOXA genes, the *F11* and *Cyp8b1* gene lost their super-enhancers (**Figure 3E**). The *Alb* gene retained its super-enhancers in *Foxa* triple null mice, but its ranking was significantly reduced (**Figure 3E**). In the absence of all three *Foxa* genes, H3K27ac enrichment was markedly decreased at the promoter and super-enhancer regions of *Alb*, *F11*, and *Cyp8b1* (**Figure 3F**). Furthermore, HNF4 α binding to the super-enhancer sites of these genes was also significantly reduced in *Foxa* triple null mice (**Figure 3F**).

To further clarify the effects of FOXA2 and FOXA3 on the formation of super-enhancers of liver function genes, we used RNAi to knock down FOXA2 alone or FOXA2 and FOXA3 simultaneously in primary human hepatocytes. ChIP-qPCR revealed that the knockdown of FOXA2 reduced the binding of H3K27ac, HNF4 α, EP300 and MED1 to the promoter and the majority of super-enhancer regions of *ALB*, *F7*, and *F10* genes (**Figure 3G**). The bindings were further reduced when hepatocytes simultaneously treated with siFOXA2 and siFOXA3 (**Figure 3G**). Silencing FOXA2 alone, or both FOXA2 and FOXA3 together, did not significantly affect the binding of super-enhancer components to the *F5* gene (**Figure 3G**). Co-immunoprecipitation (Co-IP) showed that FOXA2 directly interacts with HNF4 α but not with EP300 or MED1 (**Figure 3H**). These results suggest that, although FOXA2 does not directly interact with EP300 and Mediator, it is essential for the assembly of the EP300 and Mediator complexes at super-enhancer sites.

Consistent with the above results, ATAC-seq analysis further showed that chromatin accessibility at gene bodies and flanking regions was significantly reduced in human primary hepatocytes with FOXA2 knockdown (**Figure 3I**). Simultaneously administration of siFOXA2 and siFOXA3 further reduced chromatin accessibility in hepatocytes (**Figure 3I**). The most pronounced alteration of chromatin accessibility was observed in the regions surrounding transcription start sites (TSS), indicating widespread chromatin condensation (**Figure 3I**).

3C-qPCR assays further revealed that the interactions between promoters and super-enhancers were significantly decreased in the *ALB*, *F7*, and *F10* genes, but not in the *F5* gene, when FOXA2 was silenced by RNAi (**Figure 3J**). The interactions were further reduced in hepatocytes simultaneously treated with siFOXA2 and siFOXA3 (**Figure 3J**).

These results suggest a central role of FOXA2 in maintaining super-enhancer landscapes and chromatin accessibility in hepatocytes.

### Liver function genes form super-enhancers during LPC-to-hepatocyte differentiation

In an urgent clinical syndrome such as ALF, patients suffer from the loss of the majority of hepatocytes ^19^. In this setting, LPCs initiate the expression of vital liver function genes to maintain essential liver function ^20^. It raises an interesting question: do LPCs also possess super-enhancers on vital liver function genes?

To answer this question, we used the human LPC line HepaRG cells to establish an LPC-to-hepatocyte differentiation model according to a previous report ^21^. After incubation with DMSO for 14 days, a portion of HepaRG cells were differentiated into hepatocyte-like cells (**Figure S4A**). Periodic acid-Schiff staining showed glycogen deposition in these differentiated cells (**Figure S4A**). Immunofluorescence (IF) staining for ALB and HNF4 α confirmed the hepatocyte phenotype in these cells (**Figure S4B**). Co-IF staining further demonstrated that these HNF4 α -positive hepatocyte-like cells also expressed robust FOXA2 (**Figure S4C**).

Subsequently, we purified these hepatocytes-like cells for further investigation. RNA-seq was performed on undifferentiated LPCs (HepaRG cell), hepatocyte-like cells and human primary hepatocytes (HPHs) to investigate dynamic transcriptome alterations during LPC differentiation. Compared to HPHs, differentiated LPCs and hepatocyte-like cells demonstrated a similar transcriptome profile (**Figure S4D**). In contrast, un-differentiated LPCs exhibited lower levels of *SLC2A2*, *ADH4*, *CYP2E1*, *CYP8B1*, *F7*, *F11* and *F12* transcript compared to HPHs (**Figure S4D**). Interestingly, un-differentiated LPCs also exhibited relative high levels of *HNF4A* and *ALB* transcripts compared to differentiated LPCs (**Figure S4D**).

To investigate epigenetic changes during LPCs differentiation, we performed ATAC-seq and ChIP-seq for H3K4me1, H3K27ac, FOXA2, and FOXA3 in undifferentiated LPCs and hepatocyte-like cells. The average signal profile and heatmap displayed that overall chromatin accessibility and the enrichment of H3K4me1 were not different between undifferentiated and differentiated LPCs (**Figure 4A**). However, enrichments of H3K27ac, FOXA2, and FOXA3 were significantly higher in hepatocyte-like cells than in LPCs (**Figure 4A**). Based on the enrichment of H3K27ac signals, 185 genes with typical enhancers and 73 genes with super-enhancers were identified in undifferentiated HepaRG cells (**Figure 4B**). The 73 genes with super-enhancers did not include any of the aforementioned ten genes (e.g., *ALB*, *APOB*, *CYP8B1*, *HNF4A*, *F5*, *F7*, *F10*, *F11*, *F12*, and *SLC2A2*) (**Figure 4B**). In hepatocyte-like cells (HLCs) at day 18 and day 28, the number of typical enhancers increased to 2708 and 7025, while super enhancers increased to 194 and 454, respectively (**Figure 4B**). At day 18, the ten mentioned liver function genes acquired typical enhancers but not super-enhancers (**Figure 4B**). By day 28, the *ALB*, *APOB*, and *SLC2A2* genes had acquired super-enhancers, while the other genes exhibited high activity on their typical enhancers (**Figure 4B**). ChIP-seq analysis showed that H3K27ac, FOXA2, and FOXA3, but not H3K4me1, were more enriched in the ten liver function genes in hepatocyte-like cells compared with undifferentiated HepaRG. Compared to cells at day 18, the enrichment of H3K27ac, FOXA2, and FOXA3 was further increased in hepatocyte-like cells at day 28 (**Figure 4C** and **Figure S5A**). Consistent with the finding based on ChIP-seq, IF analysis demonstrated robust H3K27ac expression in HNF4 α -positive hepatocyte-like cells compared to surrounding undifferentiated HepaRG cells (**Figure S5B**). ATAC-seq analysis revealed increased chromatin accessibility at the promoters and super-enhancer regions of the *ALB*, *APOB*, *CYP8B1*, and *SLC2A2* genes in hepatocyte-like cells compared to undifferentiated HepaRG cells (**Figure 4C** and **Figure S5A**). In the other genes, their promoter accessibility also increased, whereas the super-enhancer regions retained similar accessibility (**Figure 4C** and **Figure S5A**). These findings suggest that certain liver function genes—such as *ALB*, *SLC2A2*, and *APOB*—establish super-enhancers at the hepatocyte-like cell stage. In contrast, other liver function genes have not yet formed super enhancers but still exhibit strong enrichment of H3K27ac, FOXA2, and FOXA3 at their promoters and/or prospective super-enhancer regions.

**Figure 4.**
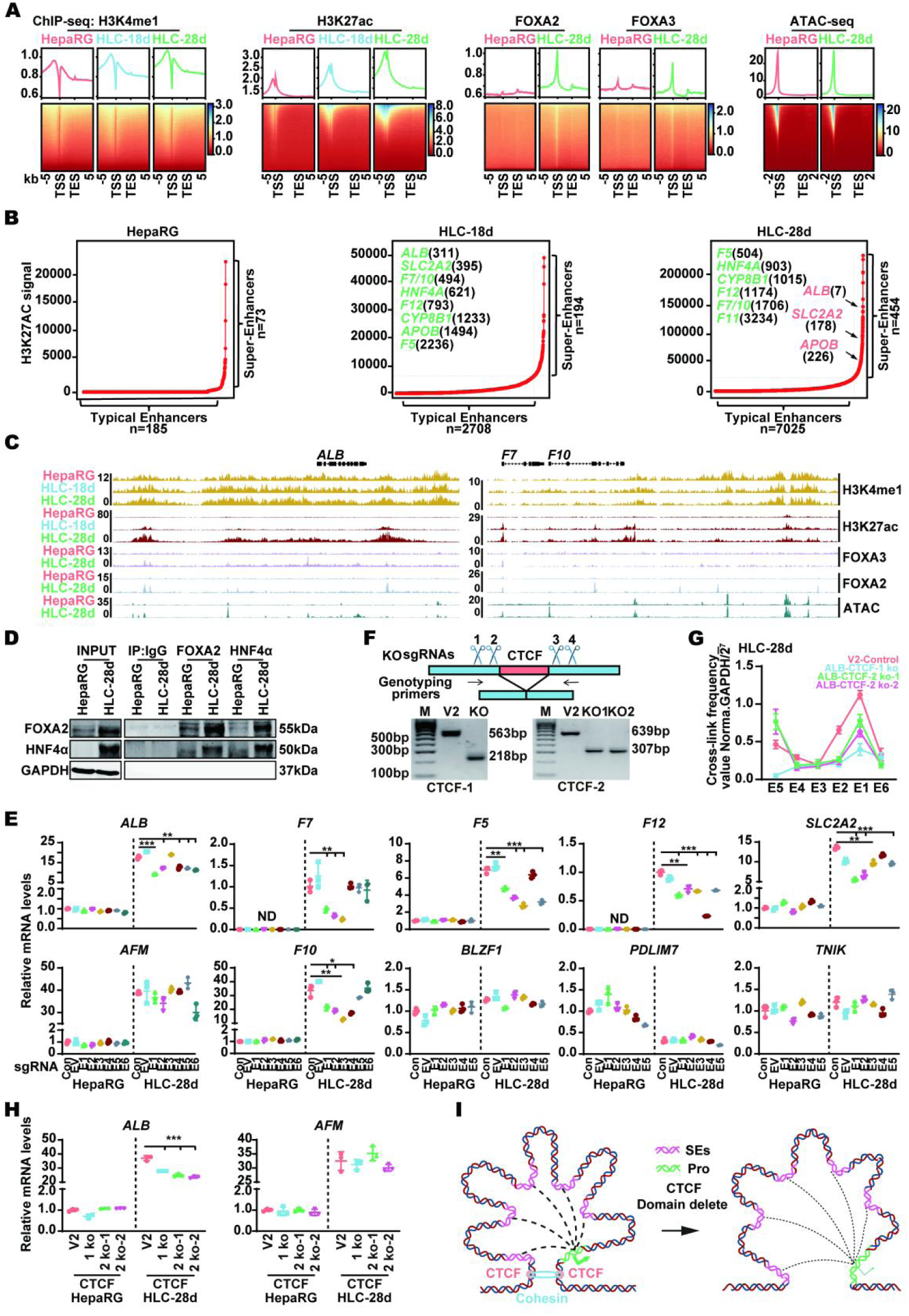
Formation of super-enhancers at liver function genes in LPCs during differentiation toward hepatocytes. (A) Normalized read distribution profiles of ChIP-seq (H3K4me1, H3K27ac, FOXA2, and FOXA3) and ATAC-seq are presented for liver progenitor cells (HepaRG) and hepatocyte-like cells (HLCs) at day 18 and day 28. (B) Typical enhancers (Green) and super-enhancers (red) were identified based on H3K27ac signals in HepaRG and hepatocyte-like cells. (C) ChIP-seq and ATAC-seq tracks highlight H3K4me1, H3K27ac, FOXA2 and FOXA3 binding, as well as chromatin accessibility at *ALB*, *F7* and *F10* loci in HepaRG and hepatocyte-like cells. (D) Co-immunoprecipitation was performed to measure the interaction between FOXA2 and HNF4α in undifferentiated HepaRG and hepatocyte-like cells. (E) Expression levels of mRNA for the indicated genes following treatment with control sgRNA (Con), empty vector (EV), or CRISPRi-mediated super-enhancers repression in LPCs and hepatocyte-like cells are shown. PPIA, measured by qRT-PCR, was used as a normalization control. Each point represents an independent biological replicate. Data are presented as mean ± standard error of the mean (SEM) from three independent experiments. P values were calculated using a two-sided Student’s t-test; *P < 0.05, **P < 0.01, ***P < 0.001. (F) Top: A schematic depicts CRISPR-knockout (KO) of CTCF binding sites. Bottom: Genotyping PCR results show homozygous knockout of two CTCF binding sites in the super-enhancers of the *ALB* gene in LPCs. (G) 3C-qPCR was used to detect the chromatin loop at the super-enhancer in hepatocyte-like cells, in which the CTCF binding sites flanking the *ALB* super-enhancers were deleted. Data are shown as mean ± SEM. Statistical significance was determined using appropriate statistical tests: *p < 0.05, **p < 0.01, ***p < 0.001. (H) qPCR was performed to examine *ALB* and *AFM* expression in HepaRG and hepatocyte-like cells where the CTCF binding sites flanking the *ALB* super-enhancers were deleted. Data are shown as mean ± SEM. Statistical significance was determined using appropriate statistical tests: *p < 0.05, **p < 0.01, ***p < 0.001. (I) A model illustrates CTCF- and Cohesin-mediated chromatin looping at super-enhancers.

Furthermore, 3C-qPCR assays revealed that the interactions between promoters and super-enhancers were significantly increased in the *ALB*, *F7*, and *F10* genes, but not in the *F5* gene (**Figure S5C**). Consistent with Co-IF staining, Co-IP analysis confirmed a significantly increased FOXA2 and HNF4α interaction in hepatocyte-like cells compared to undifferentiated HepaRG cells (**Figure 4D**).

As in human primary hepatocytes, we established CRISPR-dCas9 interference targeting the super-enhancers of the *ALB*, *F5*, *F7*, *F10*, *F12*, and *SLC2A2* genes in undifferentiated LPCs and hepatocyte-like cells. Interfering with these super-enhancers affected the mRNA expression of the examined genes only in hepatocyte-like cells, but not in un-differentiated LPCs (**Figure 4E**). With the exception of E4 in the *F5* super-enhancer, E4, E5 and E6 in the *F7* super-enhancer, E5 and E6 in the *F10* super-enhancer, and E4 in the *SLC2A2* super-enhancer, disruption of any other super-enhancer elements significantly reduced RNA expression of *ALB*, *F5*, *F7*, *F10*, *F12*, and *SLC2A2* (**Figure 4E**). Similar to hepatocytes, the CRISPR interference did not alter the mRNA levels of *AFM*, *BLZF1*, *PDLIM7*, and *TNIK*, which are located outside the TADs containing the *ALB*, *F5*, *F7*, *F10*, *F12*, and *SLC2A2* super-enhancers, respectively (**Figure 4E**). These findings suggest that putative super-enhancer regions associated with liver function genes (e.g., *F5*, *F7*, and *F10*) exert significant regulatory effects on gene expression, although they have not yet developed mature super-enhancers. These genes have the potential to develop into fully mature super-enhancers at the later stage of the LPC-to-hepatocyte differentiation.

To further clarify how super-enhancers influence gene transcription, we constructed lentiCRISPR-v2-sgRNA to delete two CTCF binding sites flanking the *ALB* super-enhancer (**Figure 4F** and **Figure S5D**). 3C-qPCR assays revealed that the interaction between multiple sites in super-enhancers and the promoter was significantly reduced upon CTCF deletion (**Figure 4G**), accompanied by a marked decrease in *ALB* mRNA level, while *AFM* expression remained unchanged (**Figure 4H**). These results illustrate the critical role of super-enhancer structure in gene transcription (**Figure 4I**).

Collectively, these results suggest that undifferentiated LPCs do not possess super-enhancers on vital liver function genes. However, these genes form super-enhancers during LPC-to-hepatocyte differentiation. Interfering with super-enhancers impairs the transcription of liver function genes.

### FOXA2 is essential for the expression of liver function genes with super-enhancers in LPCs

Next, we investigated the role of FOXA2 and FOXA3 in in regulating liver function genes with super-enhancers in LPCs. The enrichments of FOXA2 and FOXA3 within the super-enhancers of these genes significantly increased as LPC differentiated into hepatocyte-like cells (**Figure 4A-B** and **Figure S5A**). When hepatocyte-like cells were treated with siRNA-FOXA2 or siRNA-FOXA2 combined with siRNA-FOXA3, the mRNA expression of liver function genes with super-enhancers (*ALB*, *APOB*, *CYP8B1*, *F7*, *F10*, *F11*, *F12*, and *HNF4A*) was remarkably reduced (**Figure 5A**). Notably, silencing FOXA2 also reduced the mRNA expression of *ALB*, *APOB*, *F10*, *F11*, and *HNF4A* in un-differentiated HepaRG cells (**Figure 5A**).

**Figure 5.**
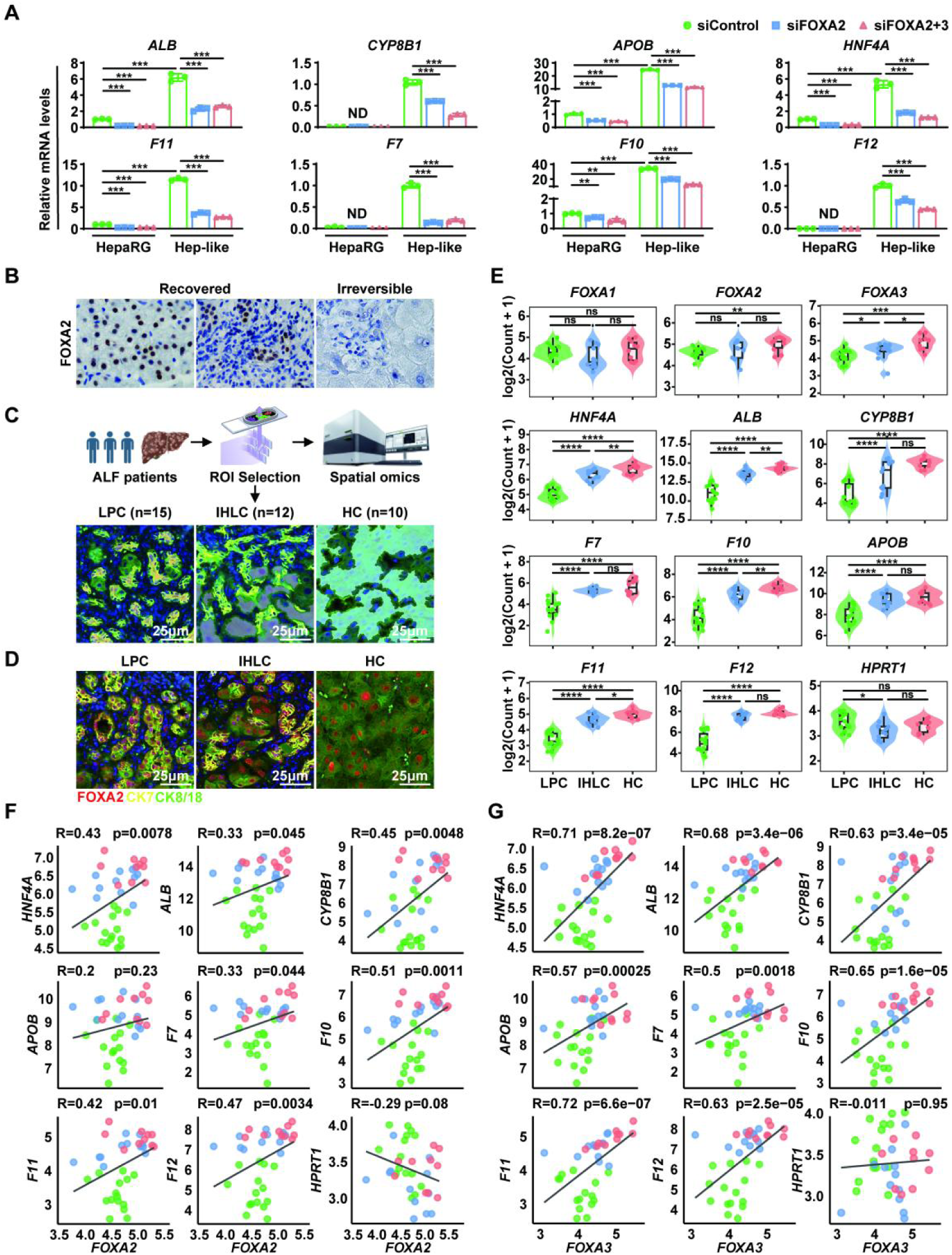
FOXA2 expression is closely associated with liver function gene expression and clinical outcome in patients with ALF. (A) qPCR was performed to examine the mRNA expression of liver function genes with super-enhancers in human primary hepatocytes treated with siRNA-FOXA2 or siRNA-FOXA2/3. *PPIA*, measured by qRT-PCR, was used as a normalization control. Each point represents an independent biological replicate. Data are presented as the mean ± standard error of the mean (SEM) from three independent experiments. P values were calculated using a two-sided Student’s t-test; ND, not detected, *P < 0.05, **P < 0.01, ***P < 0.001. (B) Immunohistochemical staining for FOXA was performed in collected liver tissues. Three representative images demonstrate FOXA2 expression in hepatocytes (**a**) and LPCs (**b**) in recovered ALF patients but not in irreversible (**c**) ALF patients. (C) A scheme depicts the GeoMx DSP workflow. A total of 37 regions of interest (ROIs) were collected based on cell-type-specific markers: CK7^+^ /Ck8/18^-^ LPCs (n = 17), CK7^+^ /Ck8/18^+^ intermediate hepatocyte-like cells (IHLC, n = 12), and CK7^-^/CK8/18^+^ hepatocytes (HC, n = 10). Gray-masked regions indicate the precise ROIs selected for transcriptomic profiling within each cell type. (D) Co-immunoprecipitation for CK7, CK8/18, and FOXA2 was performed to measure FOXA2 expression in LPCs, IHLCs, and hepatocytes in ALF patients. FOXA2 expression is shown in a representative ALF patient. (E) Base on the spatial transcriptomic expression (log2-transformed counts +1), violin plots show the expression levels of transcription factors *FOXA1*, *FOXA2*, *FOXA3*, and *HNF4A*, as well as liver function genes (*ALB*, *CYP8B1*, *APOB*, *F7*, *F10*, *F11*, *F12*) and control gene *HPRT1* in LPCs, IHLCs and HCs. ns, not significant; *p < 0.05; **p < 0.01; ***p < 0.001; ****p < 0.0001. (**F-G**) Pearson correlation was performed to assess the relationship between *FOXA2* (F) or *FOXA3* (**G**) transcripts and liver function genes with super-enhancers (*ALB*, *APOB*, *CYP8B1*, *F7*, *F10*, *F11*, *F12*, and *HNF4A*), as well as control gene *HPRT1* in liver tissues collected from ALF patients. Correlation coefficients (R) and p values are indicated for each plot.

To investigate the potential effects of FOXA2 in patients, we performed immunohistochemical (IHC) staining for FOXA2 in 20 MHN-associated ALF patients, including 5 recovered patients and 15 irreversible patients who underwent liver transplantation. These patients had been analyzed in a previous study ^22^. We found that recovered patients showed low MELD scores and robust hepatic HNF4 α expression, either in LPCs or leftover hepatocytes, whereas irreversible patients demonstrated high MELD scores and weak HNF4 α expression ^22^. IHC staining for FOXA2 revealed that robust FOXA2 expression was only detected in the recovered ALF patients (**Figure 5B**). Given that ALF patients suffer from MHN, leading to the loss of the majority of hepatocytes in some cases, recovered patients showed FOXA2 expression in LPCs, whereas irreversible patients did not (**Figure 5B**). Impressively, the remaining hepatocytes in the recovered patients also demonstrated robust FOXA2 expression (**Figure 5B**).

Next, we performed a GeoMx digital spatial transcriptomics analysis on liver tissues collected from 2 irreversible ALF patients (**Figure 5C**). IHC for CK7 and CK8/18 was used to identify LPCs (CK7-positive), intermediate hepatocyte-like cells (IHLC) (both CK7- and CK8/18-positive), and hepatocytes (CK8/18-positive) (**Figure 5C**). We collected multiple regions of interest (ROIs) from LPCs (n=15), IHLCs (n=12) and CK8/18-positive hepatocytes (n=10) **(Figure 5C)**. We also performed co-IF for FOXA2 in the four patients. FOXA2 protein expression was detected in the liver tissues undergoing spatial transcriptome analysis (**Figure 5D**). Based on the spatial transcriptome analysis, we analyzed transcripts of FOXA genes and the previously mentioned liver function genes with super-enhancers (*ALB*, *APOB*, *CYP8B1*, *F7*, *F10*, *F11*, *F12*, and *HNF4A*) and *HPRT1* gene as a control.

FPKM analysis revealed that the levels of transcripts for *FOXA2*, *FOXA3*, *ALB*, *APOB*, *CYP8B1*, *F7*, *F10*, *F11*, *F12*, and *HNF4A* genes gradually increased from LPCs to IHLCs and hepatocytes, but not *FOXA1* (**Figure 5E**). Pearson correlation analyses further showed robust correlations between *FOXA2/FOXA3* and liver function genes with super-enhancers (p<0.01 for all correlations, **Figure 5F-G**).

Finally, we analyzed the associations between *FOXA2, FOXA3*, and liver function genes associated with super-enhancers in two publicly available RNA-seq databases (GSE38941 and GSE14668), which included liver tissues collected from healthy individuals and HBV-ALF patients. Compared to healthy controls, two cohorts of HBV-associated ALF patients demonstrated significantly reduced transcript levels of FOXA2, FOXA3, *ALB*, *APOB*, *CYP8B1*, *F7*, *F10*, *F11*, *F12* and *HNF4A* (**Figure S6A-B**). In both databases, Pearson correlation analysis revealed a strong positive correlation between *FOXA2* or *FOXA3* transcript levels and the examined genes associated with super-enhancers (p<0.01 for all correlations, **Figure S6C-D**). These results illustrate that FOXA2 and FOXA3 may play a critical role in regulating the expression of these liver function genes in ALF patients.

## Discussion

Over the last decade, the concept of the super-enhancer has emerged, highlightinh their critical roles in various physiological and pathophysiological processes, such as embryonic stem cell and carcinogenesis ^4^. Although there are different opinions regarding the nomenclature of “super-enhancer”, their pivotal functions in cell identity, embryonic development and cancer pathogenesis are widely recognized ^23^. Young and colleagues have generated a catalog of super-enhancers in 86 human cells and tissues types ^4^, illustrating their widespread presence in human cells. Given the unusually high levels of transcriptional activity associated with super-enhancers, we hypothesized that essential liver function genes in hepatocytes may be regulated by such epigenetic structures, enabling these cells to efficiently produce liver function proteins in response to physiological or pathophysiological challenges.

This study confirms the presence of super-enhancers in the majority of liver functional genes in hepatocytes. In both human and mouse livers, more than 1,000 genes possess super-enhancers, encompassing most liver function genes (**Table S1** and **S2**). Furthermore, applying CRISPR-dCas9 interference to target the super-enhancers or using lentiCRISPR-v2-sgRNA to delete two CTCF binding sites flanking the super-enhancers significantly inhibits the expression of vital liver function genes, such as *ALB* and coagulation factors, in hepatocytes. These results underscore the essential role of super-enhancers in ensuring the robust expression of critical liver function genes in hepatocytes.

In MHN-associated ALF, where a severely damaged liver has lost most or all hepatocytes, LPCs take over hepatocyte functions to provide essential liver proteins that support systemic homeostasis ^7–9^. A key question is whether LPCs also possess super-enhancers in vital liver function genes, similar to hepatocytes. We found that quiescent LPCs do not form super-enhancers in liver function genes. However, during the differentiation of LPCs into hepatocytes, super-enhancers in these genes are established. In an *in vitro* model using the human LPC line HepaRG, quiescent LPCs were found to have only 93 genes with super-enhancers, none of which are liver function genes. After 18 and 28 days of culture, the number of super enhancers in these cells increased to 194 and 454, respectively. Vital liver function genes, including *ALB*, *APOB*, and *CYP8B1* were identified with super-enhancers by day 28. Notably, the current LPC differentiation model spans two stages over 28 days: the first 14 days are dedicated to proliferation, followed by 14 days of differentiation. Both day 18 and day 28 are part of the proliferation phase, suggesting that the formation of super-enhancers in liver function genes occurs during the early phase of LPC activation. These findings partially explain clinical observations, where LPCs robustly express *HNF4A*, *ALB* and coagulation factors in recovered ALF patients ^22, 24^.

How do hepatocytes maintain super-enhancers at liver function genes, whereas LPCs form super-enhancers at these genes only during activation? Lessons from embryonic stem cells (ESCs) provide a clue to the answer: In ESCs, cell-type specific master transcription factors play a key role in the formation of super-enhancers ^14^. For example, Oct4, Sox2 and Nanog are master transcription factors that regulate the formation of super-enhancers at most genes controlling the pluripotent state by recruiting the Mediator complex and maintaining chromatin accessible in ESCs ^14^. During embryonic liver development, six transcription factors—HNF4 α, HNF1 α, HNF1β, HNF6, LRH-1, and FOXA—form a core transcription network that governs the differentiation of hepatoblasts into hepatocytes ^25–27^. Among these, LRH-1 and FOXA2 are pioneer factors, which have the capacity to maintain chromatin accessible ^27, 28^. Joo and colleagues reported that LRH-1 is a key regulator of a super-enhancer-associated network in hepatocytes ^5^. In APAP-treated mice, LRH-1 was shown to regulate 3% of super-enhancer-associated genes ^5^. In the present study, we investigated the role of FOXA2 in regulating super-enhancer-associated liver function genes. Our findings reveal that super-enhancers at many liver function genes demonstrate FOXA2 occupancy along with other enhancer-associated components (e.g., H3K4me1, EP300, and MED1) and hepatic master transcription factor HNF4α. Compared to normal hepatocytes, which possess 1,332 super-enhancers, the number of super-enhancers decreased to 1,154 in mouse hepatocytes isolated from *Foxa2*-deficient mice. ChIP-qPCR analysis further showed that knocking down FOXA2 significantly reduces the binding of H3K27ac, HNF4 α, EP300 and MED1 to the promoters and the majority of super-enhancer regions of *ALB, F7* and *F10* genes, indicating that FOXA2 is required for the assembly of the EP300 and Mediator complexes at super-enhancer regions. However, subsequent Co-IP analysis revealed that FOXA2 directly interacts with HNF4α but not with EP300 or MED1. How FOXA2 affects the assembly of the EP300 and Mediator complexes remains to be elucidated.

In addition to FOXA2, FOXA1 and FOXA3 are also enriched within the super-enhancers of multiple liver function genes. Subsequent experiments using individual or combined silencing of FOXA factors in hepatocytes and mice showed interaction between FOXA1, FOXA2, and FOXA3: FOXA1 antagonizes, while FOXA3 enhances, FOXA2’s role in regulating target gene transcription. The collaboration between FOXA2 and FOXA3 is critical for maintaining chromatin accessibility at gene bodies and flanking regions. Simultaneous administration of siFOXA2 and siFOXA3 further reduced chromatin accessibility compared to silencing FOXA2 alone. Notably, the impact of FOXA2 and FOXA3 on chromatin accessibility is more predominant in the transcription start site regions compared to super-enhancer sites, suggesting that FOXA2 and FOXA3 regulate gene transcription by modulating both promoter and enhancers. This mechanism was further confirmed by 3C-qPCR analysis: the interactions between promoters and super-enhancers were significantly reduced in multiple liver function genes when FOXA2 was silenced by RNAi.

In the absence of *Foxa2*, mouse hepatocytes lose nearly 200 super-enhancers while maintaining more than 1,100, indicating the existence of both *Foxa2*-dependent and *Foxa2*-independent transcription regulation. In this study, we carefully analyzed ten vital liver function genes (*ALB*, *APOB*, *CYP8B1*, *F5*, *F7*, *F10*, *F11*, *F12*, *HNF4A*, and *SLC2A2*) associated with super-enhancers. Among these genes, *ALB*, *APOB*, *CYP8B1*, *F7*, *F10*, *F11*, *F12*, and *HNF4A* are FOXA2-dependent, while *F5* and *SLC2A2* are FOXA2-independent. Based on experiments using *Foxa2*-deficient mice and loss-of-function assays in vitro, FOXA2-associated genes are primarily involved in molecular functions related to RNAPII-specific DNA-binding transcription factor binding, heparin binding, heme binding, alcohol dehydrogenase (NAD+) activity, and lipid transporter activity. These findings partially explain the critical role of FOXA2 in regulating multiple liver function genes, including coagulation factor, *ALB*, and detoxification enzymes. FOXA2-dependent expression of these liver function genes is essential in LPCs in MHN-associated ALF. Several lines of evidence highlight the critical role of FOXA2 in ALF patients: (1) In a cohort of ALF patients, only survivors exhibit robust hepatic FOXA2 expression; (2) FOXA2 expression is closely associated with MELD scores; (3) In a database of HBV-associated ALF patients, strong positive correlations are observed between FOXA2 expression and vital liver function genes, including *ALB*, *APOB*, *CYP8B1*, *F7*, *F10*, *F11*, *F12*, and *HNF4A*. (4) Spatial transcriptome analysis further demonstrates a close correlation between FOXA2 and these liver function genes in LPCs. Given FOXA2 forms a complex with HNF4 α (**Figure 3G**), it is unsurprising that *FOXA2* expression is highly associated with *HNF4A*. Specifically, (1) robust expression of both factors is observed only in survivors; and (2) in survivors, the protein expression of FOXA2 and HNF4α in LPCs was localized to areas without hepatocytes, while remaining hepatocytes still exhibit high levels of FOXA2 and HNF4 α ^22^. These findings not only elucidate the mechanisms by which master hepatic transcription factors, such as FOXA2 and HNF4 α, control vital liver function in ALF but also provide real translational significance in clinical practice. Can FOXA2 and HNF4α serve as prognostic markers in ALF patients? Could ALF treatment be enhanced by upregulating the expression of these two factors? These important translational questions require further investigation.

Collectively, super-enhancers represent a critical epigenetic feature that enables ample transcription of liver function genes in response to both normal and pathophysiological demands, owing to their unusually high levels of enhancer activity. In urgent clinical syndromes such as ALF, the formation of super-enhancers in vital liver function genes is essential for LPCs to take over liver function and sustain the patients’ life. FOXA2 is a key regulator in maintaining super-enhancers in liver function genes in both hepatocytes and LPCs. In addition to regulating super-enhancers, FOXA2 collaborates with HNF4 α to control the transcription of numerous vital liver function gene in both cell types. The expression of these transcription factors is closely associated with clinical outcomes in MHN-associated ALF. Maintaining the expression of these two master hepatic transcription factors may represent a viable therapeutic strategy in clinical practice.

One of the unanswered questions in this study is how activated LPCs form super-enhancers at liver function genes. Although FOXA2 is critical for maintaining super-enhancers, FOXA2 itself is not sufficient to drive super-enhancer formation at liver function genes in LPCs. The main evidence for this comes from our *in vitro* LPC differentiation model, where RNA-seq and qPCR analyses revealed high levels of FOXA2 expression in undifferentiated LPCs. However, these LPCs do not form super-enhancers at liver functional genes. Brown et al. reported that TNF-α can induce super-enhancer formation by activating NF-κB in endothelial cells ^29^. When LPCs were stimulated with TNF-α, they failed to form super-enhancers at liver functional genes. Furthermore, TNF-α reduced FOXA2 expression in LPCs (data not shown). Thus, the mechanism underlying super-enhancer formation at liver function genes in activated LPCs requires further investigation.

## Materials and Methods

### Patients

Twenty patients with ALF were enrolled from the Department of Medicine II, University Medical Center Mannheim, Medical Faculty Mannheim, Heidelberg University and the Department of Gastroenterology and Hepatology, Beijing You’an Hospital, Affiliated with Capital Medical University ^22^. Liver tissues were collected when patients received tumor excision or liver transplantation. ALF is defined as a severe liver injury, leading to coagulation abnormality usually with an international normalized ratio ≥ 1.5, and any degree of mental alteration (encephalopathy) in a patient without pre-existing liver disease or with an illness of up to 4 weeks duration ^8^. The study protocol was approved by local ethics committees (Jing-2015-084, and 2017-584N-MA). Written informed consent was obtained from the patients or their representatives.

### Cells

Human primary hepatocytes (HPHs) were isolated by the Cell Isolation Core Facility of the Biobank Großhadern, University Hospital, LMU Munich. Mouse primary hepatocytes (MPHs) were isolated from *ad libitum* fed mice using the two-step collagenase perfusion technique. The isolated primary hepatocytes were seeded on collagen-coated plates in Williams’ E medium supplemented with 100 nM dexamethasone, 5% Insulin-Transferrin-Selenium, 10% (v/v) heat-inactivated FBS, 1% L-glutamine and 100 U/ml penicillin G/streptomycin sulfate. The isolation of primary hepatocytes has been described previously ^30^.

Undifferentiated HepaRG cells (HPR101), purchased from Biopredic International (Saint Gregoire, France), were cultured in Williams E medium supplemented with 10% (v/v) heat inactivated FBS, 5 µg/ml insulin, 50 µM hydrocortisone hemisuccinate, 1% L-glutamine and 100 U/ml penicillin G/streptomycin sulfate. According to a previous report ^21^, undifferentiated HepaRG were initially seeded at a density of 2×10^4^ cells/cm^2^ and culture for 14 days to allow proliferation. To induce cell differentiation, 1.5% DMSO were added into the media. The cells were cultured for additional 14 days, during which they differentiated into hepatocyte-like cells (**Figure S4**). Subsequently, the hepatocyte-like cells were isolated from remaining un-differentiated HepaRG cells using mild trypsinization. The eluted cells were further separated by centrifugation at 1500g for 5 minutes on a 30% OptiPrep gradient, followed by a second centrifugation on a 15% OptiPrep gradient, to obtain pure hepatocyte-like cells.

HEK293T cells, used for lentiviral construction, were grown in DMEM medium supplemented with 10% FBS, 1% L-glutamine, and 100 U/ml penicillin G/streptomycin sulfate. All cells were cultured at 37°C in a humidified atmosphere with 5% CO_2_.

### Animals

*Foxa2*^flox/flox^ C57BL/6JGpt mice (8-10 weeks old) were obtained from GemPharmatech Co., Ltd. (Nanjing, China). Hepatocyte-specific *Foxa2* knockout (*Foxa2*^ΔHep^) mice were generated by crossing *Foxa2*^flox/flox^ mice with Alb-iCre transgenic mice. All animals were housed in the animal research facility of Anhui Medical University under standard conditions (22 °C, 12-hour light/dark cycle) with free access to food and water. All procedures were approved by the Animal Experiment Ethics Committee of the First Affiliated Hospital of Anhui Medical University and conducted in accordance with the ARRIVE (Animal Research: Reporting of In Vivo Experiments) guidelines.

### Immunohistochemistry

Immunohistochemical (IHC) staining was performed as description previously ^24^. Briefly, the sections were deparaffinized using a series of ethanol dilutions, followed by three washes with phosphate-buffered saline (PBS). Subsequently, the sections were transferred to 10 mM sodium citrate buffer (pH 6.0) for antigen unmasking in a microwave. After cooling, the sections were incubated in peroxidase blocking reagent (Dako) for 1 hour. The primary antibody was added, and the sections were incubated overnight at 4°C. The next day, EnVision peroxidase (Dako) was applied for 1 hour at room temperature following a PBS wash. The sections were developed with diaminobenzidine for 5 minutes and counterstained with hematoxylin (Merk). The dehydrated sections were mounted using malinol mounting medium (Sigma-Aldrich) for imaging.

### siRNA transfection

All siRNAs used in this study were purchased from Dharmacon and were transfected into cells with Lipofectamine RNAiMAX (Invitrogen, USA) according to the manufacturer’s instruction. Cells were subjected to different treatments following 48h transfection. Detailed information of siRNA are listed in Supplementary data.

### RNA extraction, real time-qPCR and RNA sequencing

Total RNA was extracted from cells by TRIzol™ (Thermo Fischer Scientific, USA) according to the manufacturer’s instructions. Reverse transcription was performed to synthesis cDNA using RevertAid H Minus Reverse Transcriptase (Thermo Fischer Scientific, USA). RNA quality was assessed with the Agilent 2100 Bioanalyzer and the RNA 6000 Nano Kit (Agilent, Waldbronn). The qRT-PCR assays were performed using POWRUP SYBR MASTER MIX (Thermo Fischer Scientific, USA) on a StepOnePlus Real-time PCR instrument (Applied Biosystems, USA). Samples with RNA integrity number above 9.5 were used for RNA sequencing. RNA sequencing was conducted by BGI Tech Solutions Co. (Hong Kong, China).

### RNA-seq data processing

Quality analysis of the raw FASTQ files was performed using the FastQC software. Adapter removal and low-quality sequence filtering were conducted using Trimmomatic software. Gene expression quantification in the transcriptome was achieved by aligning the cleaned RNA sequencing data to the human reference genome using the HISAT2 software. FeatureCounts software was used to count reads, accurately, and gene expression level was quantified using the FPKM or TPM calculation methods. To identify differentially expressed genes, the “limma” R package was employed, with a significance threshold set at an adjusted P-value < 0.05 and a log2-transformed fold change (log2FC) > 0.58 (FC > 1.5).

### ChIP-qPCR and ChIP-seq

ChIP was performed using the SimpleChIP® Enzymatic Chromatin IP Kit. Briefly, cells were cross-linked with 1% formaldehyde for 10 minutes at room temperature, quenched with 5 M glycine for 5 minutes at room temperature, and then washed three times with PBS. Chromatin was prepared and incubated with micrococcal nuclease for 20 minutes at 37°C, followed by sonication to a length of approximately 150-500 bp. Chromatin was immunoprecipitated by incubation with 5 µg of anti-H3K4me1, H3K27ac, 10 µg of FOXA2, FOXA3, EP300, MED1 or rabbit IgG overnight at 4°C. The next day, chromatin and antibodies were incubated with magnetic beads for 4 hours at 4°C with rotation, followed by three washes with low salt wash buffer and one wash with high salt wash buffer at 4°C for 5 minutes each. Chromatin was eluted with ChIP elution buffer at 65°C for 30 minutes with gentle vortexing and then treated with 5 M NaCl and proteinase K at 65°C overnight to reverse crosslinks. The samples were subsequently incubated with RNase at 37°C for 1 hour. ChIP DNA was purified and then detected by qPCR or sequencing. ChIP sequencing was performed by BGI Tech Solutions Co. (Hong Kong, China).

### ATAC-seq

ATAC-seq was performed using the Zymo-Seq ATAC Library Kit (Zymo Research). Briefly, 50,000 live cells were harvested and lysed on ice in 25 µL of ATAC-S buffer and 25 μL of ATAC-Lysis buffer for 5 minutes, followed by the addition of 1 mL of cold ATAC Wash Buffer and immediate centrifugation at 1000 × g for 10 minutes at 4°C. The nuclei pellets were resuspended in 50 μL of transposition buffer and 2.75 μL Tn5 enzyme, then incubated at 37 °C for 30 minutes. The purified transposed DNA was subjected to library amplification using the ATAC Library PCR Mix and UDI Tag Primer, followed by paired-end sequencing (2 × 150 bp) on an Illumina NovaSeq 6000.

### ChIP-seq and ATAC-seq data processing

Quality control of the raw FASTQ files was performed using the fastp software. Reads were aligned to the reference genome using the Bowtie2 software. Post-alignment processing was carried out using Samtools software, and formats were manipulated and duplicates marked using Picard software. Coverage files were generated, scores calculated, and reads shifted using DeepTools software. Peak identification was performed using MACS3 software, signal differences were quantified using Manorm software, and peaks were annotated using HOMER software. Bias correction, footprint score calculation, and motif detection were conducted using TOBIAS software. Finally, super enhancer ranking analysis was carried out using ROSE software.

### Identification of Super-enhancers

H3K27ac peaks were used to define enhancer boundaries, excluding those overlapping ENCODE blacklisted regions and RefSeq-annotated gene promoter regions. The remaining H3K27ac peaks were identified as enhancers and subsequently processed using the ROSE (Ranking of Super Enhancers) algorithm ^14^.

### Enhancer-Promoter Interaction Analysis Using ChIA-PET Data

CTCF- and RNA Polymerase II (RNAPII)-mediated chromatin interaction data for HepG2 cells were obtained from the ENCODE project (accession numbers: ENCSR411IVB for CTCF ChIA-PET and ENCSR857MYZ for RNAPII ChIA-PET). Raw sequencing data were processed using the ENCODE ChIA-PET pipeline to identify high-confidence chromatin loops. Topologically associating domains (TADs) encompassing super-enhancers were delineated based on combined analysis of CTCF ChIA-PET loops, CTCF ChIP-seq, and Hi-C contact maps.

Enhancer-gene associations were established by identifying the frequencies of chromatin interactions linking H3K27ac- and RNAPII-enriched enhancer regions to gene promoters. These interactions were visualized alongside Hi-C contact matrices using the WashU Epigenome Browser.

### 3C-qPCR assay

The 3C-qPCR assay was performed following the protocol described by Hagège et al^31^. In brief, human primary hepatocytes were cross-linked with 2% formaldehyde for 10 minutes at room temperature. The cross-linking reaction was quenched with 1M glycine for 5 minutes. The cells were lysed in a Lysis Buffer containing 10 mM Tris-HCl (pH 7.5), 10 mM NaCl, 5 mM MgCl2, 0.1 mM EGTA, and 1x complete protease inhibitor for 10 minutes. Nuclear pellets were obtained by centrifugation at 400 g for 5 minutes and resuspended in 1.2x restriction enzyme buffer with 0.3% SDS for 1 hour at 37°C. Subsequently, the chromatin was treated with 2% Triton X-100 for 1 hour at 37°C. The chromatin was digested overnight at 37°C using the BglII (*ALB*, *F5*) or HindIII (*F7*, *F10*) restriction enzyme and ligated overnight with T4 DNA ligase at 16°C. Following ligation, the samples were treated with proteinase K overnight at 65°C to reverse the cross-links. RNA was removed using RNase A treatment. The 3C DNA was then purified with phenol–chloroform.

For the 3C-qPCR analysis, primers for both control and target fragments were designed to anneal within 50-150 bp upstream of the HindIII or BglII restriction sites, ensuring that all primers were oriented in the same direction. The relative quantification of the 3C-ligation products was performed using SYBR Green-based qPCR, with GAPDH primers serving as an internal negative control for normalization.

### CRISPR-mediated super-enhancer functional analysis

The function of super-enhancer elements was investigated using the CRISPR-dCas9-mediated interference (CRISPRi) system and lentiCRISPR v2 ^32, 33^. Briefly, sequence-specific sgRNAs for site-specific interference of genomic targets were designed using the CRISPR filter tool (http://crispor.gi.ucsc.edu/), and sequences were selected to minimize off-target effects. The sgRNA of CRISPR-dCas9 was selected at the center of the enhancer peak to inhibit enhancer activity. The sgRNA of CRISPR-Cas9 is selected at the two boundaries of the peak to facilitate the cutting of the entire region. The oligonucleotides were annealed using T4 polynucleotide kinase and cloned into the pLV-hU6-sgRNA-hUbC-dCas9-KRAB-Puro (Addgene ID: 71236) and the lentiCRISPR v2 vector (Addgene ID: 52961) using the BsmBI-v2 Golden Gate Assembly Kit.

For lentivirus production, HEK293T cells were co-transfected with the appropriate dCas9-KRAB, psPAX2, and pMD2.G plasmids, or with lentiCRISPR v2, psPAX2, and pCMV-VSV-G plasmids, using FuGENE®6 Transfection Reagent. The transfection medium was replaced 8 hours later with fresh medium. The lentivirus-containing medium was harvested 48 hours post-transfection, filtered through a sterile 0.45 μm filter, centrifuged at 30,000g for 6 hours, and stored at −80 °C until use.

For CRISPRi, primary hepatocytes and HepaRG cells were transduced with either lentivirus-sgControl (Con), lentivirus lacking sgRNA (empty virus, EV), or lentiviral-sgRNA targeting enhancers at a multiplicity of infection (MOI) of 30 for primary hepatocytes and an MOI of 5 for HepaRG cells. 5 μM polybrene was added to the lentiviral medium to enhance transduction efficiency. After 2 days of transduction, cells were subjected to puromycin selection: 5 μg/ml for primary hepatocytes and 1 μg/ml for HepaRG cells, for 2 days. Positively transduced HepaRG cells were expanded and used in subsequent differentiation and gene expression assays. Gene expression analysis was also performed on the selected primary hepatocytes.

For CRISPR-Cas9-mediated CTCF-binding deletion, HepaRG cells were transduced with either lentivirus-V2 (empty vector control) or a pool of lentivirus containing four different sgRNA pairs targeting the CTCF-binding sites at a multiplicity of infection (MOI) of 5. Following transduction, cells were selected with 1 μg/mL puromycin for 2–3 days and plated into 96-well plates to isolate single-cell-derived clones. Clones were screened for CRISPR-mediated deletion of the CTCF-binding regions by PCR amplification, DNA gel electrophoresis, and Sanger sequencing. To avoid genomic deletions from perturbing adjacent cis-regulatory elements, CTCF binding sites were deleted in minimal fragments (<350 bp). Validated clones containing specific deletions were amplified and used for differentiation experiments, gene expression analysis, and 3C-qPCR. All sgRNA sequences and genotyping PCR primers are listed in Table S3.

### Spatial transcriptome analysis

Formalin-Fixed Paraffin-Embedded (FFPE) tissue samples obtained from patients with ALF were analyzed using the GeoMx digital spatial profiler (NanoString) with the whole transcriptome atlas panel. Section (5 μm) were deparaffinized in xylene, rehydrated through graded ethanol, and subjected to antigen retrieval in Tris-EDTA buffer (pH 9.0, 00-4956-58; Thermo Fisher Scientific) for 20 minutes. Proteolytic digestion was performed by incubating the samples with 1 μg/mL proteinase K (AM2546; Thermo Fisher Scientific) for 15 minutes at 37°C. Following re-fixation in 10% neutral buffered formalin (15740-04; EMS Diasum) for 5 minutes, slides were hybridized overnight at 37°C with ∼18000 probes. After stringent washes, slides were stained with morphology markers: CK8/18-AF488 (hepatocytes), CK7-AF594 (LPCs) and SYTO83-CY3 (nuclei) for 1 hour. Based on the fluorescence signals, 37 regions of interests (ROIs) were defined and categorized into three cell populations: CK7^+^ /Ck8/18^-^ LPCs (n = 15), CK7^+^ /CK8/18^+^ intermediate hepatocyte-like cells (IHLC, n = 12), and CK7^-^/CK8/18^+^ hepatocytes (HC, n = 10). UV light was used to release oligonucleotide barcodes from selected ROIs, which were then PCR-amplified and purified with AMPure XP beads (A63880; Beckman Coulter) purification. The purified libraries were sequenced on the NextSeq 550 platform (Illumina) for spatial gene expression analysis.

### GeoMx data analysis

Fastq data were converted to DCC files using the GeoMx NGS Pipeline software (Version 3.1.1.6). Normalization and quality control were performed analyzed using the GeoMx DSP Data Center Software (Version 3.1.0.222). ROIs were evaluated for PCR contamination and sequencing depth. All 18,676 gene-associated probes and ROIs passed quality control. Gene expression values were nomalized using third quartile (Q3) normalization. Bioinformatic analysis were conducted in R (version 4.5.0) with the following packages: ggplot2 (v3.5.2) for data visualization, ggpubr (v0.6.0) for significance annotation and correlation statistics, dplyr (v1.1.4) for data manipulation, and svglite (v2.1.3) for vector graphic export. Base R packages, including stats and grDevices, were used for correlation testing and graphical output, respectively. Correlation between transcription factors and liver function genes was assessed using Spearman’s rank correlation. P-values were adjusted using the Benjamini-Hochberg method, with adjusted P < 0.05 considered statistically significant.

### Western Blotting

An immunoblot assay was performed as previously described. Cell protein lysates were extracted using RIPA buffer or IP buffer with protease inhibitor. Samples (30µg each) were separated by 8%-15% SDS-PAGE and then transferred to PVDF membranes. Following incubation with 5% bovine serum albumin (BSA) for 1h at room temperature, the membranes were incubated with primary antibodies overnight at 4°C. The next day, the membranes were probed with secondary antibodies and developed using a chemiluminescent substrate.

### Protein co-immunoprecipitation

Cells were lysed with 1 ml of ice-cold Pierce™IP Lysis Buffer (with protease inhibitors). Clarified protein lysates were incubated with 5 μg of the indicated primary antibodies or control IgG overnight at 4°C and then added to Protein A/G Magnetic Beads for 2 h at 4°C with rotation. The beads were washed four times with lysis buffer, resuspended in 40 μL of 2×SDS sample buffer, and heated at 95°C for 10 minutes to elute bound proteins. The samples were subsequently analyzed by Western blotting as described above.

### Immunofluorescence staining

The cells were fixed on slides with 4% paraformaldehyde (PFA) for 10 minutes at room temperature, followed by three washes with PBST (PBS containing 0.1% Tween-20). Permeabilization was performed using 0.25% Triton X-100 in PBST for 5 minutes. After additional PBST washes, the cells were blocked with 5% bovine serum *ALB* (BSA) in PBST for 1 hour at room temperature. Subsequently, the slides were incubated with primary antibodies diluted in buffer overnight at 4°C. The next day, the slides were washed three times with PBST and incubated with Alexa Fluor®-conjugated secondary antibodies (1µg/mL) for 1 hour at room temperature in the dark. After three additional washes, the nuclei were counterstained with DRAQ5 for 5 minutes. Images were captured using a Leica TCS SPE confocal microscope.

### Periodic Acid-Schiff Staining

Glycogen (stained purple) was detected using a periodic acid–Schiff (PAS) staining kit. Briefly, the cells were fixed with 4% paraformaldehyde and treated with 0.5% periodic acid for 10 minutes at room temperature. After being rinsed with distilled water, the samples were incubated with Schiff’s reagent for 15 minutes in the dark. The slides were then washed with running tap water for 5-10 minutes. The nuclei were counterstained with hematoxylin, and images were captured using a light microscope.

### Data Availability

ChIP-and ATAC seq data of HepG2 cells were obtained from ENCODE: HNF4 α (ENCSR469FBY), FOXA1 (ENCSR865RXA), FOXA2 (ENCSR066EBK), FOXA3 (ENCSR092OVN), CTCF (ENCSR607XFI), EP300 (ENCSR271XMW), MED1 (ENCSR959XNY), RAD21 (ENCSR000BLS), H3K4me1 (ENCSR000APV), H3K27ac (ENCSR000AMO), and ATAC (ENCSR042AWH). ChIP-seq and ATAC-seq data of human livers were obtained from ENCODE: HNF4 α (ENCSR601OGE), FOXA1 (ENCSR735KEY), FOXA2 (ENCSR310NYI), CTCF (ENCSR254YRM), RAD21 (ENCSR917QNE), H3K4me1 (ENCSR554XXQ and ENCSR218ZMU), H3K27ac (ENCSR119XNK and ENCSR458RRZ), and ATAC (ENCSR837OOH). ChIP-seq data of mouse liver-1 were obtained from GSE157452. ChIP-seq data of mouse liver-2 were obtained from ENCODE: H3K4me1 (ENCSR000CAO) and H3K27ac (ENCSR000CDH). RNA-seq and ChIP-seq data of *Foxa1*/*a2*^lox/lox^, *Foxa1*/*a2*^lox/lox^ Cre_Alfp, *Foxa3*-null, and *Foxa* triple-null mice were obtained from GSE124384 and GSE140423.

### Statistical analysis

The analyses were performed using SPSS Statistics 23.0. An unpaired Student’s t-test was used to determine statistically significant differences between two groups. *P* values less than 0.05 were considered significant and represented graphically as follows: *, *P*<0.05; **, *P*<0.01; and ***, *P*<0.001.

## Acknowledgements

We thank the Human Tissue and Cell Research Foundation, a nonprofit foundation regulated by German civil law, which facilitates research with human tissue through the provision of an ethical and legal framework for prospective sample collection. We acknowledge the support of the LIMA Live Cell Imaging at Microscopy Core Facility Platform Mannheim (CFPM).

## Author’s contributions

H.L.W. conceived the project. T.L., L.Z., H.W., and H.L.W. designed experiments. T.L. performed in vitro experiments. C.T. performed spatial transcriptome analysis. C.T., H.C., and H.W. performed bioinformatic analyses. W.X., S.W., W.Z., R.L., and Y.L. undertook animal experiment. H.L., C.S., and H.D. collected the patient samples and performed pathological evaluation. S.M. and H.N. provided primary human hepatocytes. T.L. and H.L.W. drafted the article. T.L., H.C., C.T., W.X., S.W., C.M., R.L., M.P.A.E., S.D., H.D., L.Z., H.W., and H.L.W. discussed the data and edited the article critically.

## Supplementary data

### Key resource table

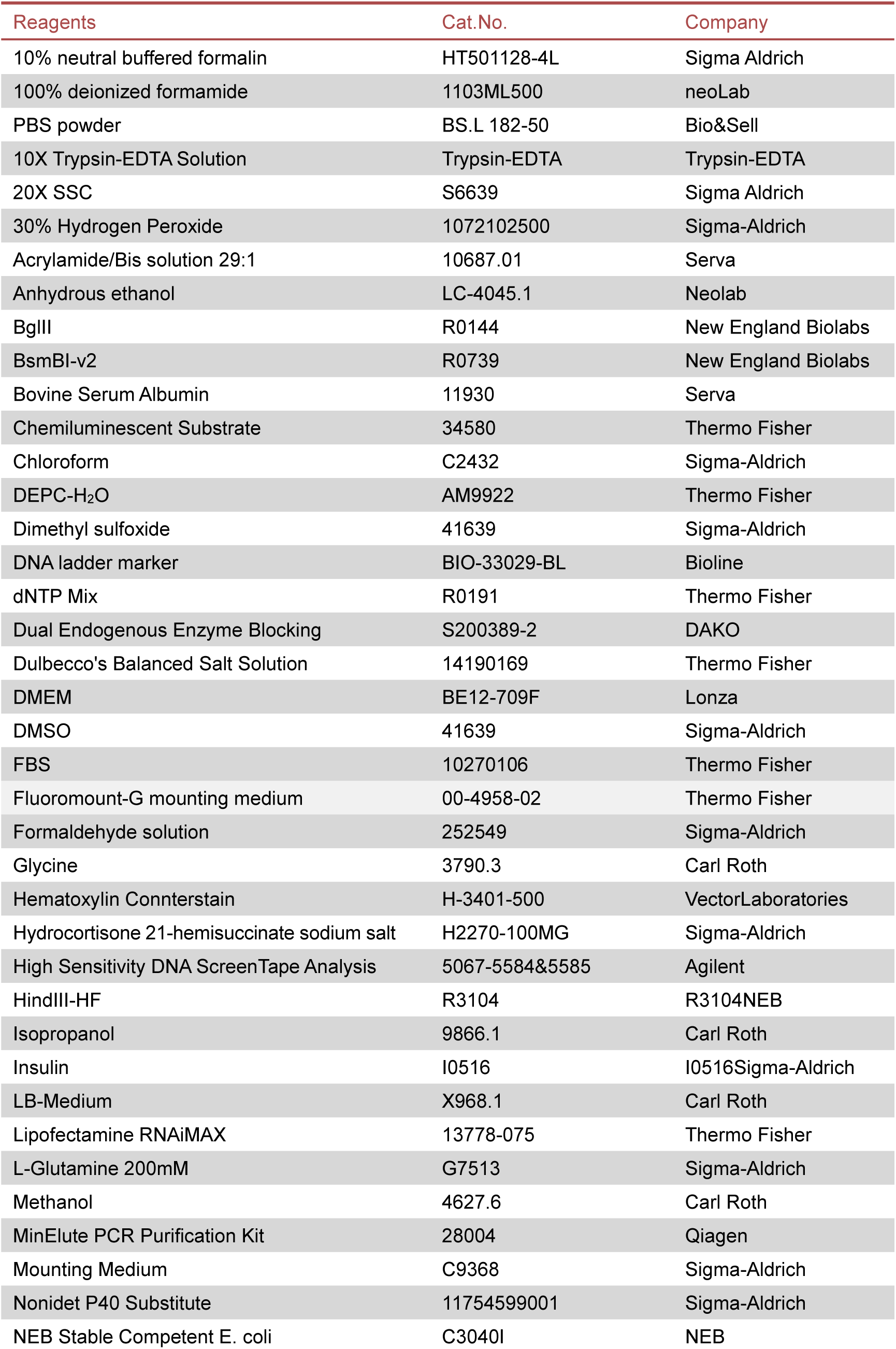
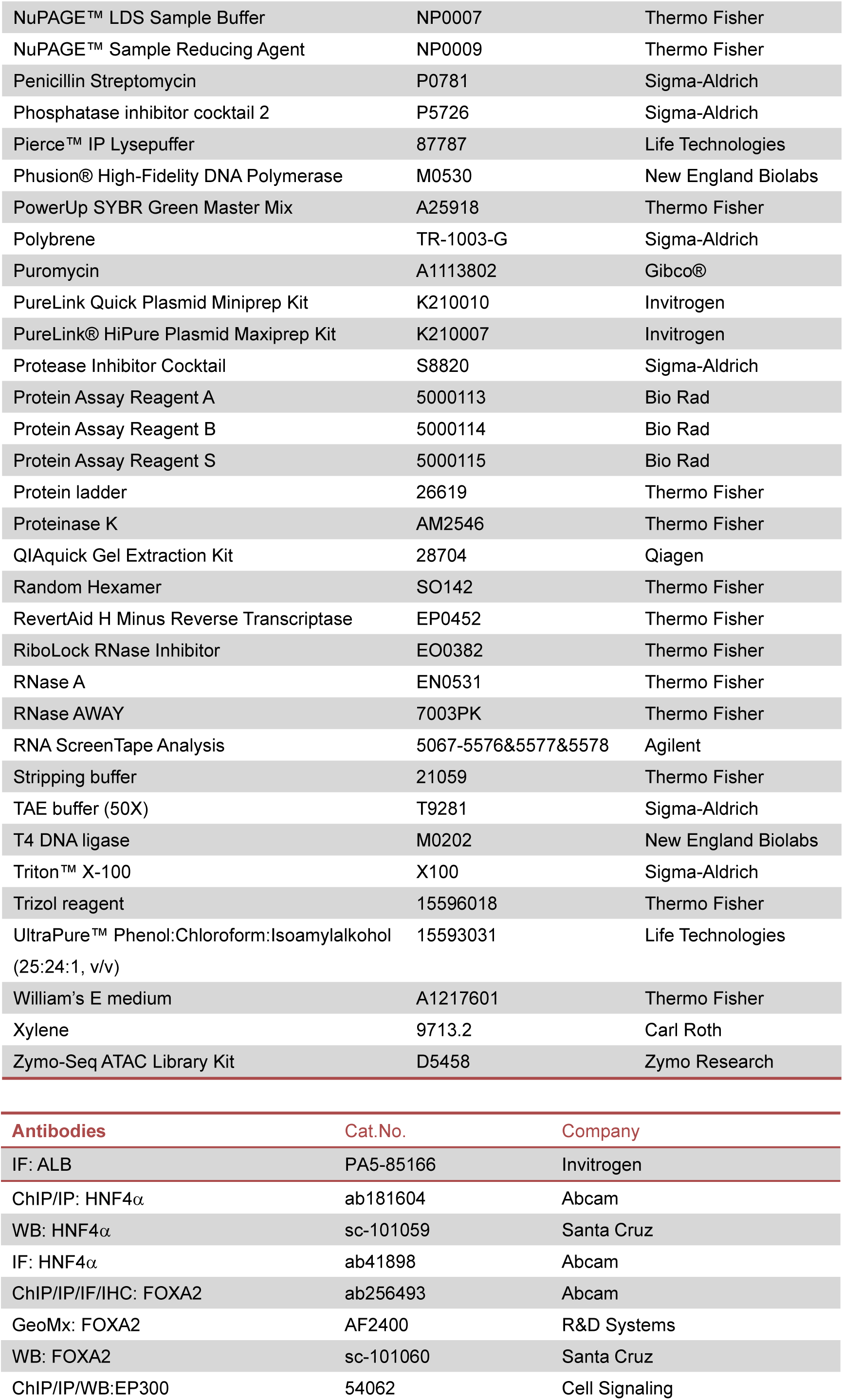
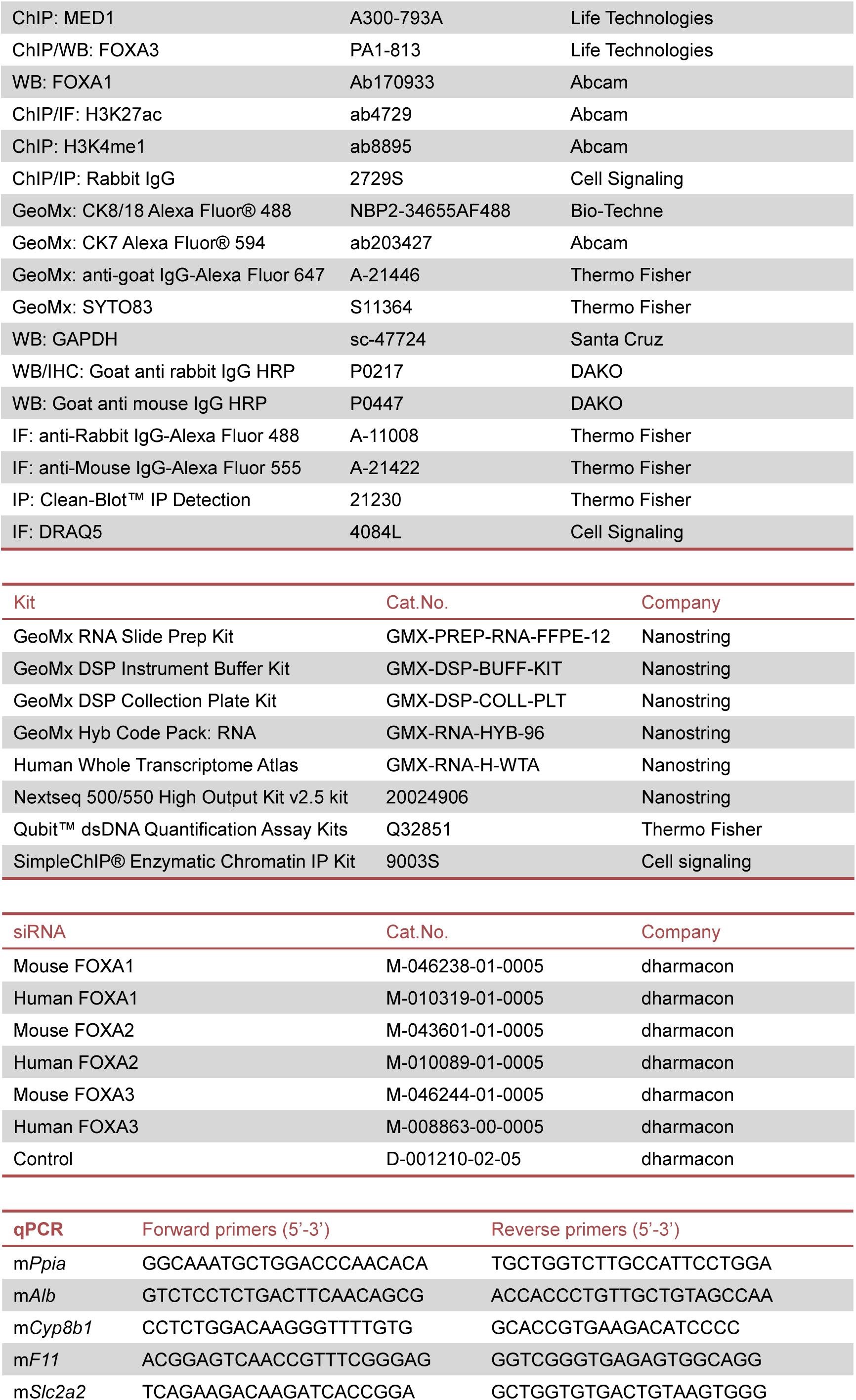
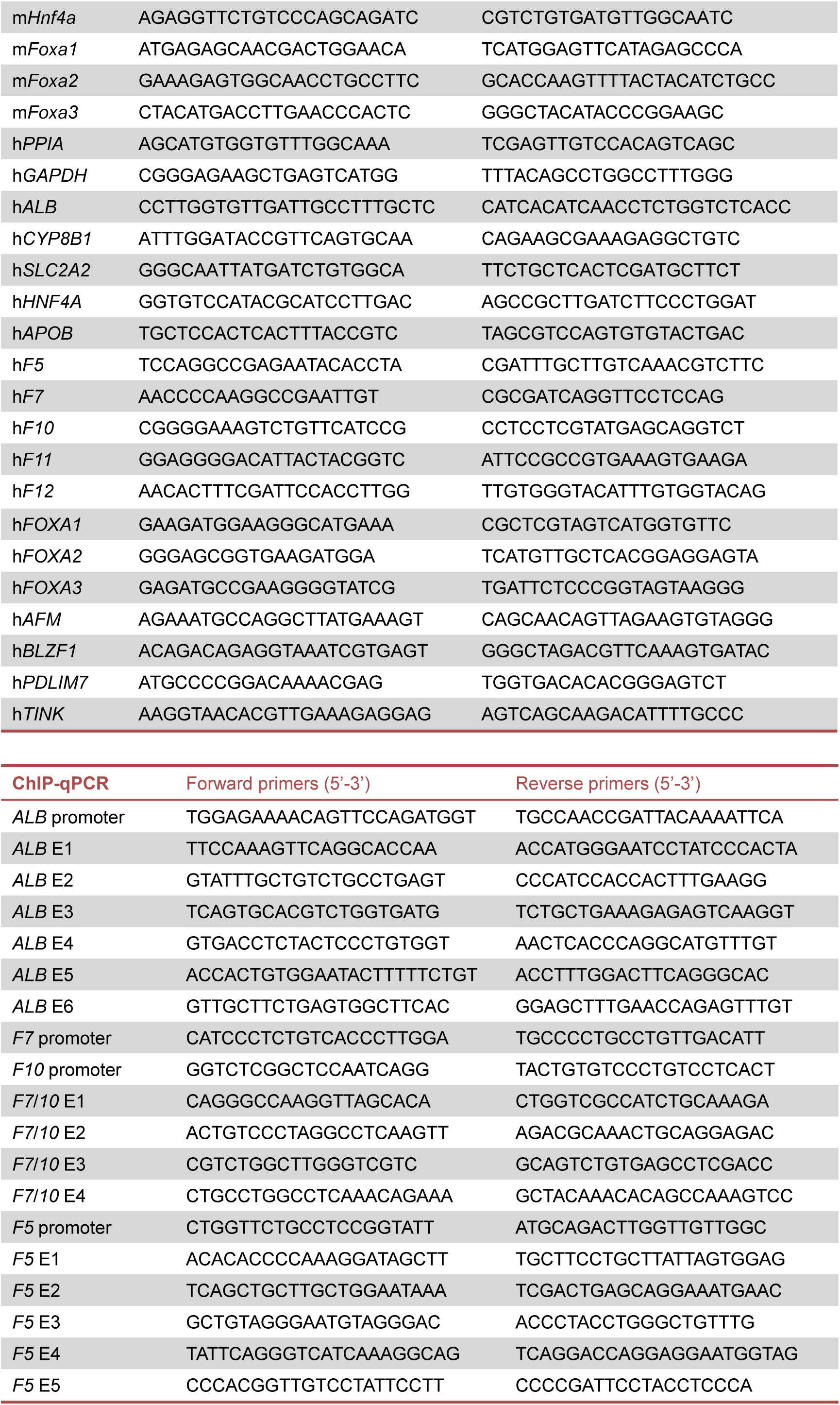
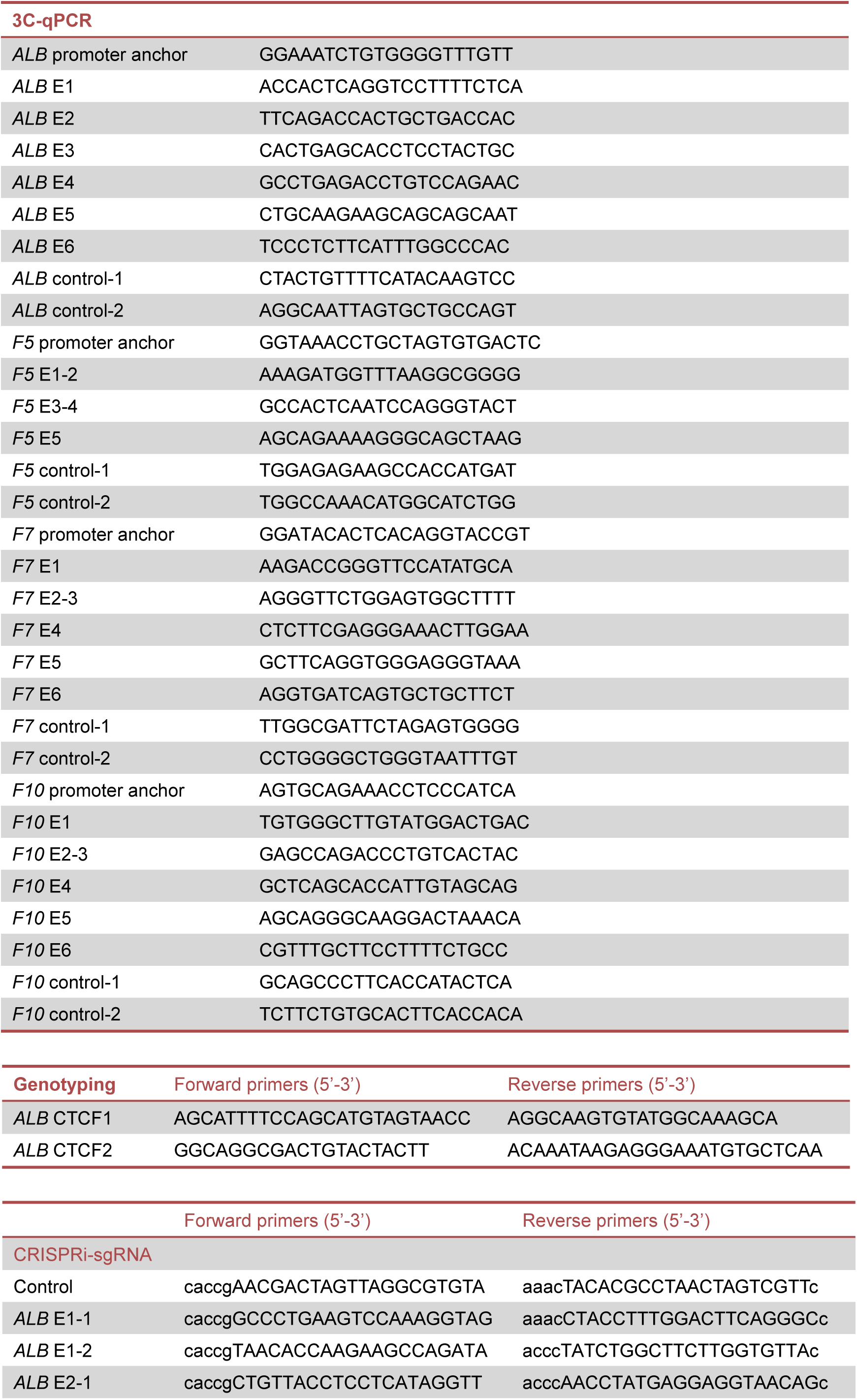
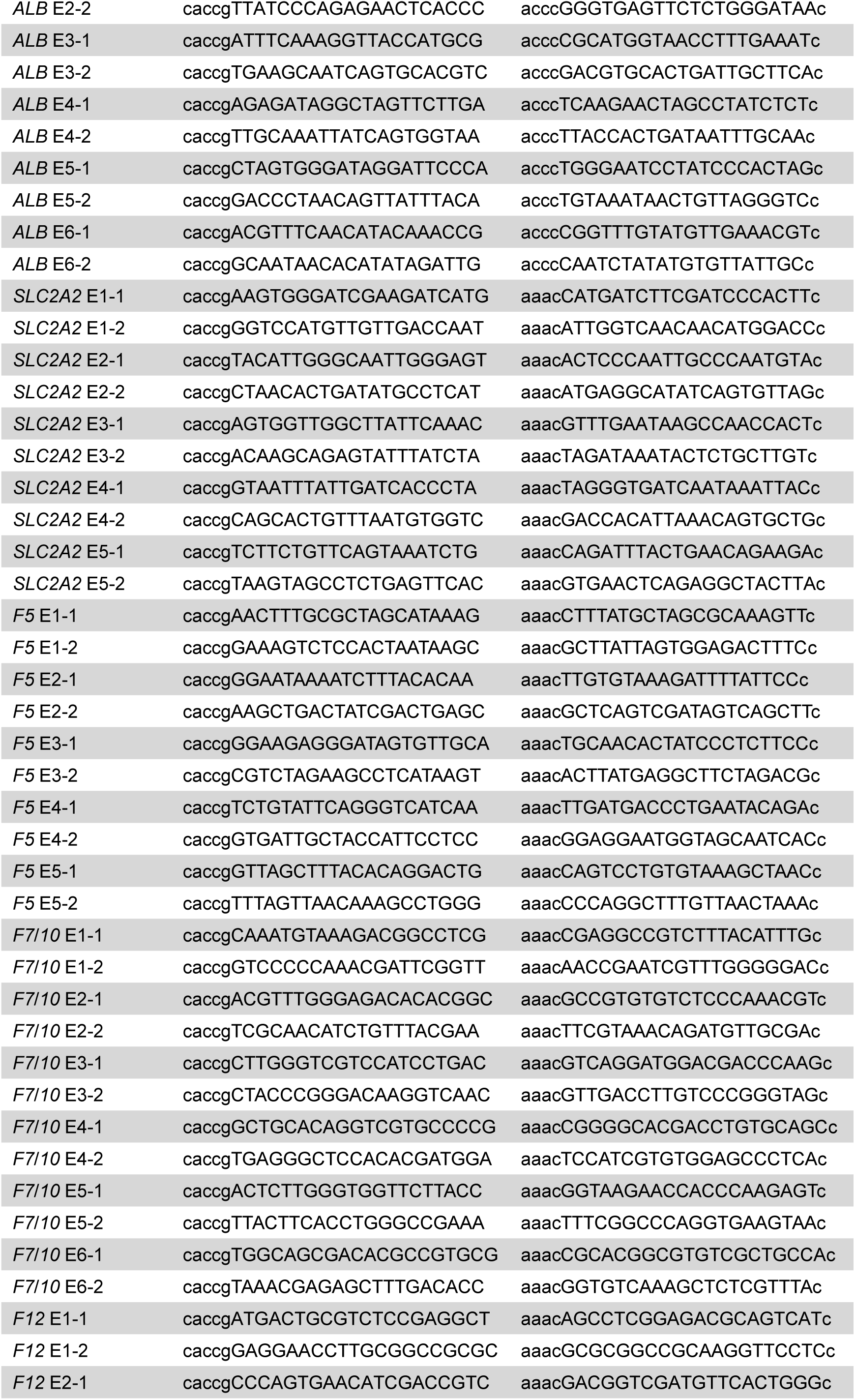
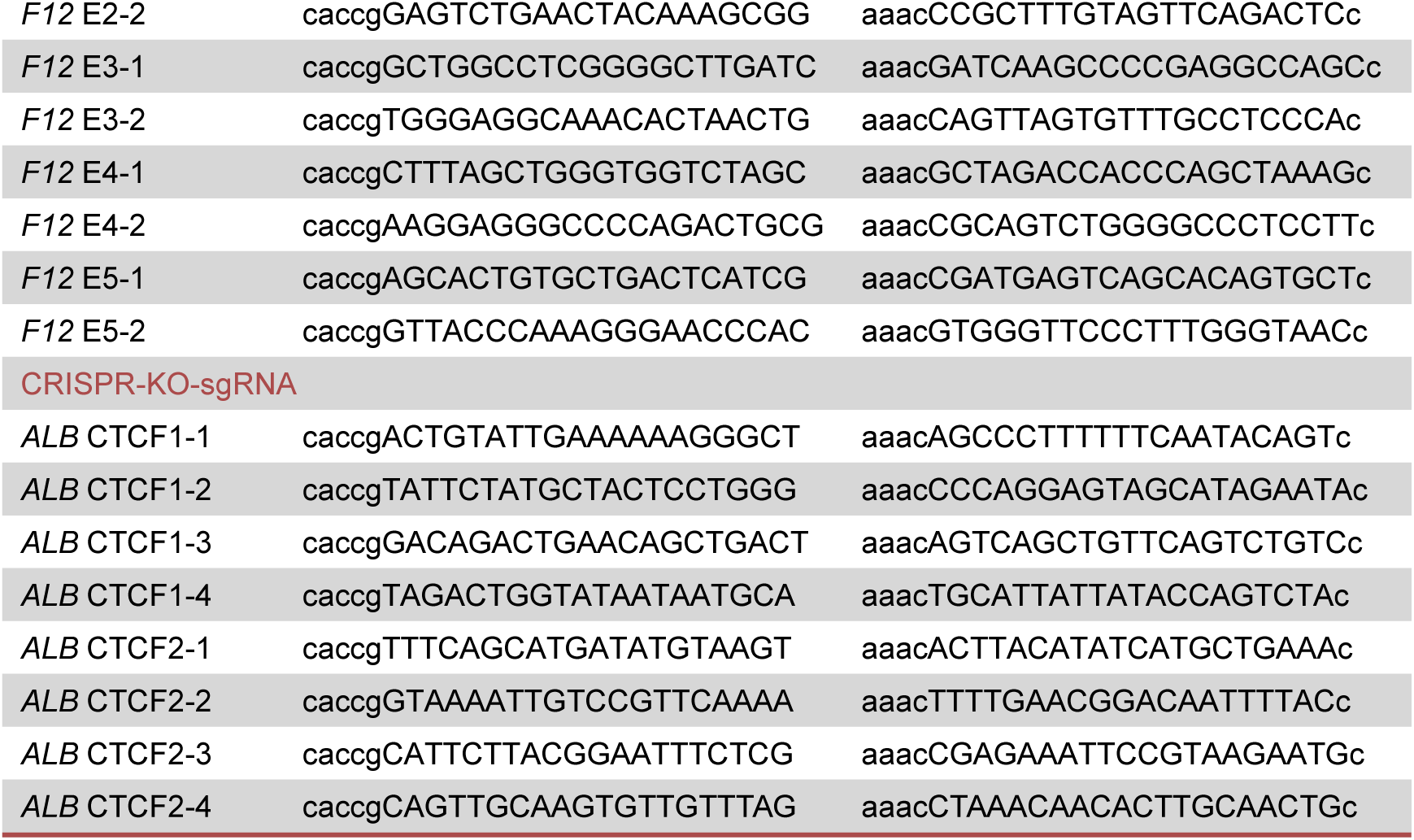

---

**Figure S1.**
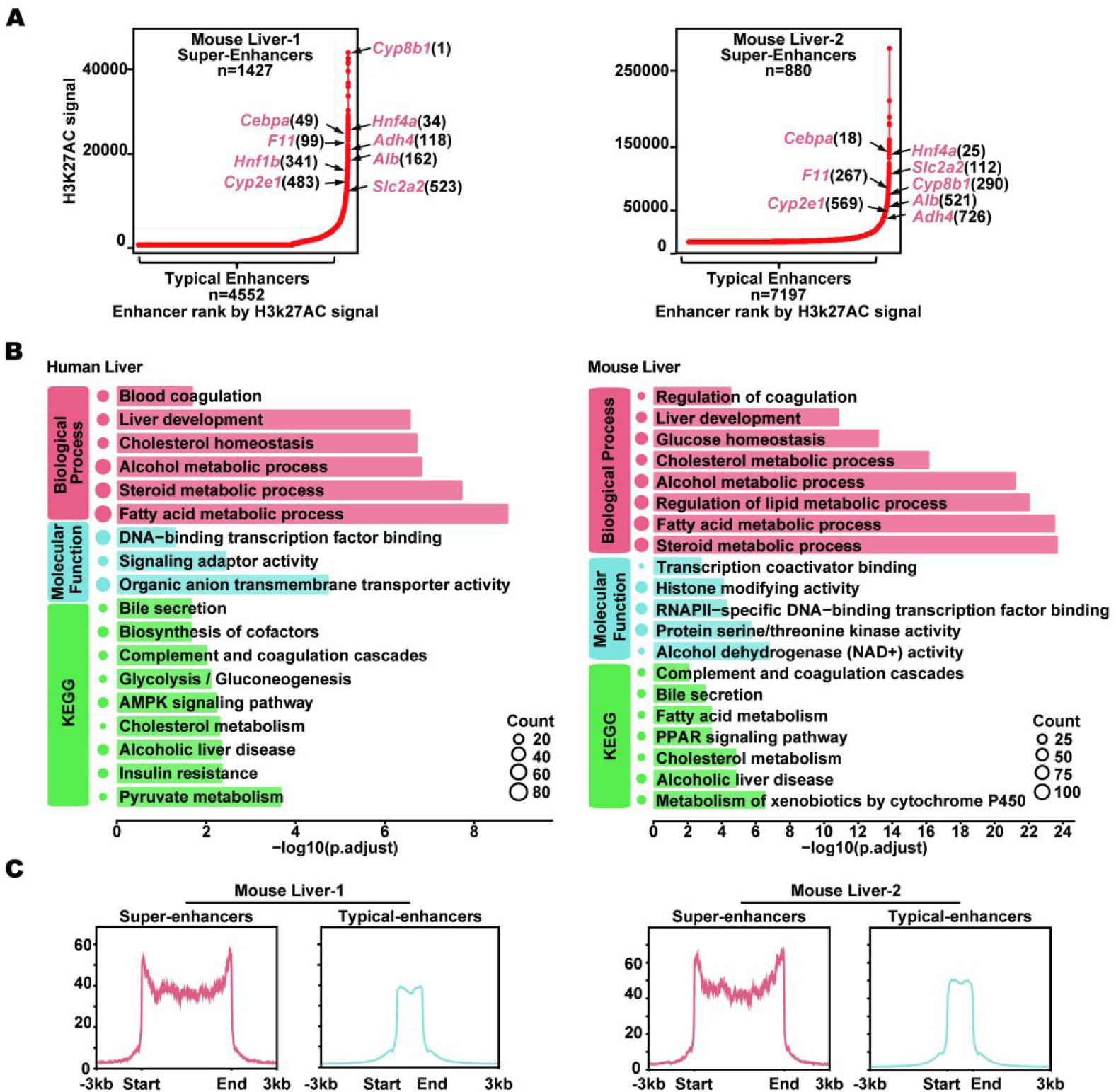
Identification of super-enhancers in human and mouse liver. **(A)** Super-enhancers were identified based on normalized H3K27ac ChIP-seq signals using the ROSE algorithm in two independent mouse liver samples. Key liver genes with SEs are marked in red with their enhancer rank. **(B)** Gene Ontology and KEGG were conducted to identify biological process and molecular function pathways associated with gene with SEs in the human and mouse liver. Bar length represents the −log10 adjusted p-value, and bubble size reflects the number of associated genes. **(C)** Average H3K27ac ChIP-seq signal is centered around SEs and TEs in mouse livers. The signal is plotted from 3 kb upstream to 3 kb downstream of the enhancer regions.

**Figure S2.**
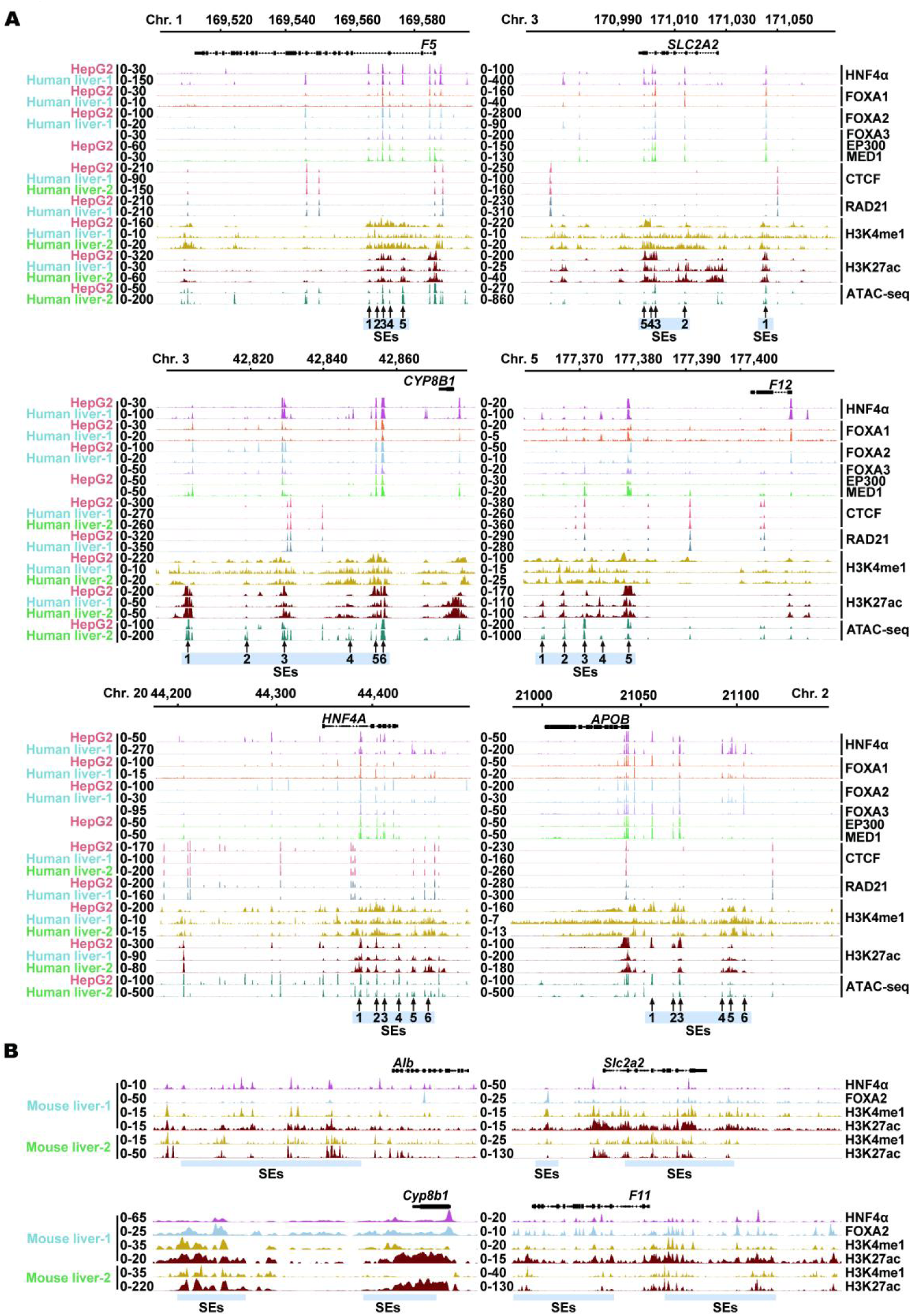
Super-enhancer landscapes at key liver function genes in human and mouse livers. (A) Genomic browser tracks show the distribution of transcription factors (HNF4 α, FOXA1, FOXA2, FOXA3), co-activators (EP300, MED1), architectural proteins (CTCF, RAD21), histone modifications (H3K4me1, H3K27ac), and chromatin accessibility (ATAC-seq) at representative super-enhancers associated loci (*F5*, *SLC2A2*, *CYP8B1*, *F12*, *HNF4A*, *APOB*) in HepG2 cells (red) and human liver tissues (blue and green). Arrows denote identified SE regions, and shaded boxes highlight regions of super-enhancers and promoters. (B) Super-enhancer profiles are shown for representative liver function genes (*Alb*, *Slc2a2*, *Cyp8b1*, and *F11*) in mouse liver samples. ChIP-seq data for HNF4α, FOXA2, H3K4me1, and H3K27ac are shown. Shaded boxes highlight regions of super-enhancers.

**Figure S3.**
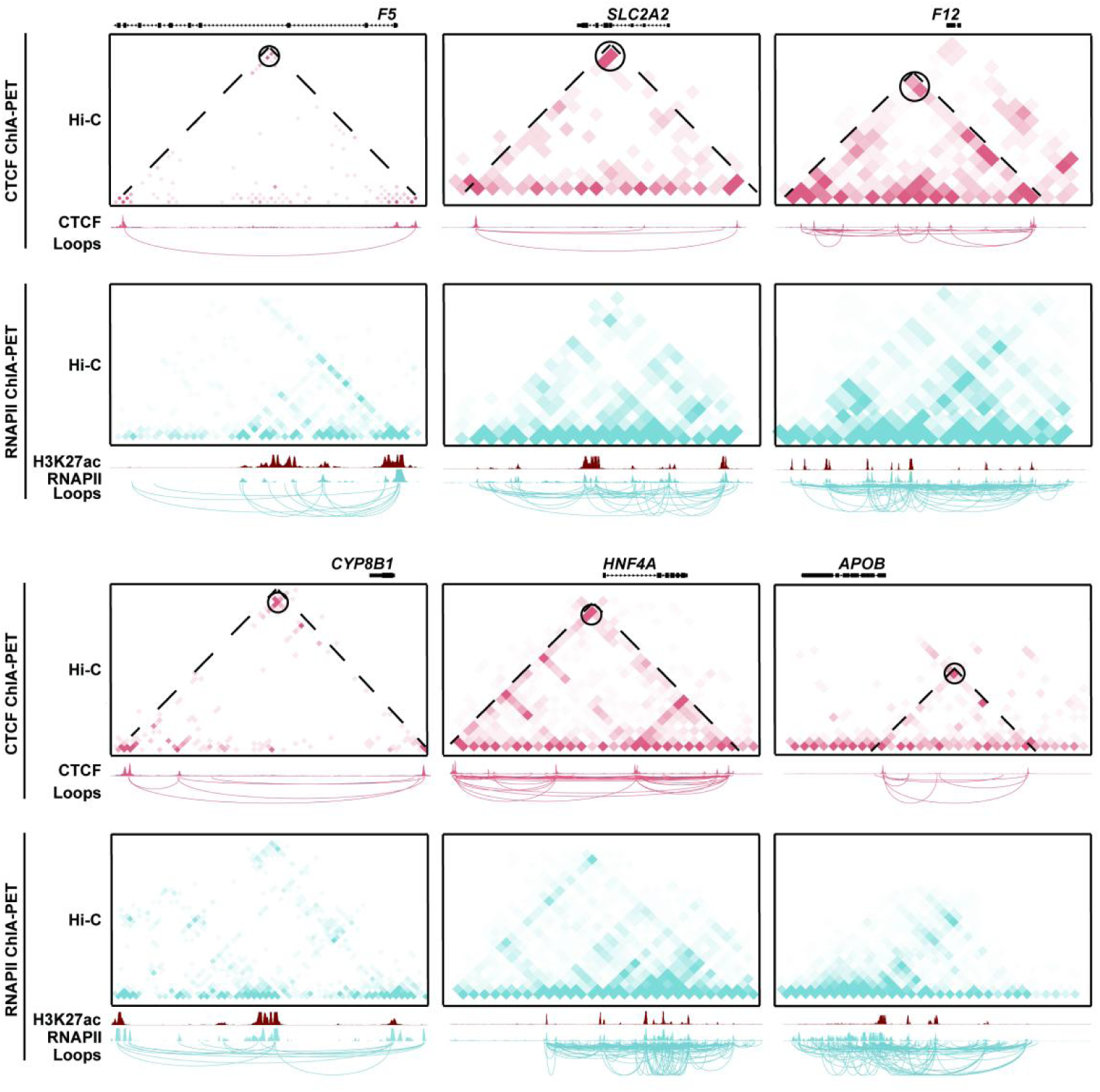
Chromatin looping at super-enhancers associated liver function genes revealed by ChIA-PET analysis. Hi-C and ChIA-PET interaction maps show domain chromatin interactions in the *F5*, *SLC2A2*, *F12*, *CYP8B1*, *HNF4A*, and *APOB* loci in HepG2 cells. Top (CTCF ChIA-PET): CTCF-mediated chromatin loops are shown in red. Hi-C contact maps reveal local topologically associated domains, while CTCF ChIA-PET loops illustrate the structural organization and long-range interactions of these regions. Bottom (RNAPII ChIA-PET): RNAPII-mediated enhancer-promoter loops, showing in blue, illustrate regulatory connections. H3K27ac ChIP-seq tracks (red) indicate active enhancer regions. RNAPII ChIA-PET maps reveal regulatory loops that connect super-enhancers to their target gene promoters.

**Figure S4.**
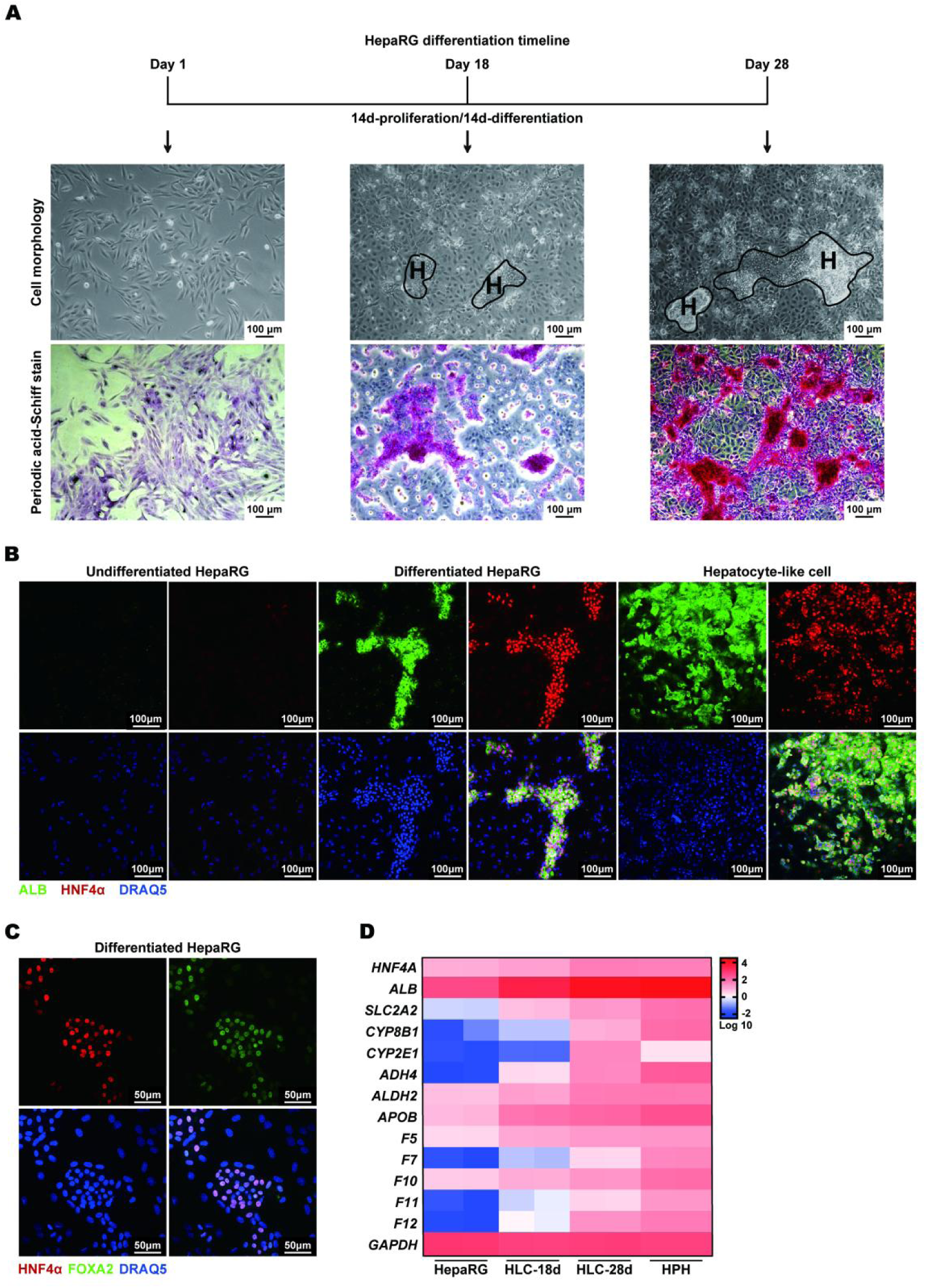
Establishment of liver progenitor cell-to-hepatocyte model in vitro. (A) A schematic depicts the timeline of HepaRG differentiation into hepatocyte-like cells. Cells were cultured for 28 days, with the first 14 days dedicated to proliferation followed by an additional 14 days for differentiation. Representative bright-field images show cell morphology at Day 1, Day 18, and Day 28. Periodic acid–Schiff (PAS) staining(pink) was performed to examine glycogen storage in those cells. H: Hepatocyte-like cells. (B) Immunofluorescence staining for ALB (Green), HNF4α (red), and nuclei (DRAQ5, blue) was performed to assess hepatocyte phenotypes in undifferentiated HepaRG cells, differentiated HepaRG cells and purified hepatocyte-like cells. (C) Immunofluorescence staining for FOXA2 (Green), HNF4 α (red), and nuclei (DRAQ5, blue) was performed in HepaRG cells (day-28). (D) Gene expression profiling during differentiation was analyzed using RNA-seq. A heatmap displays the log10-transformed expression levels of liver-specific genes in different cell types: HepaRG cells (Undifferentiated LPC), purified hepatocyte-like cells on day-18 (HLC-18d), HLCs on day-28 (HLC-28d), and primary human hepatocytes (HPH).

**Figure S5.**
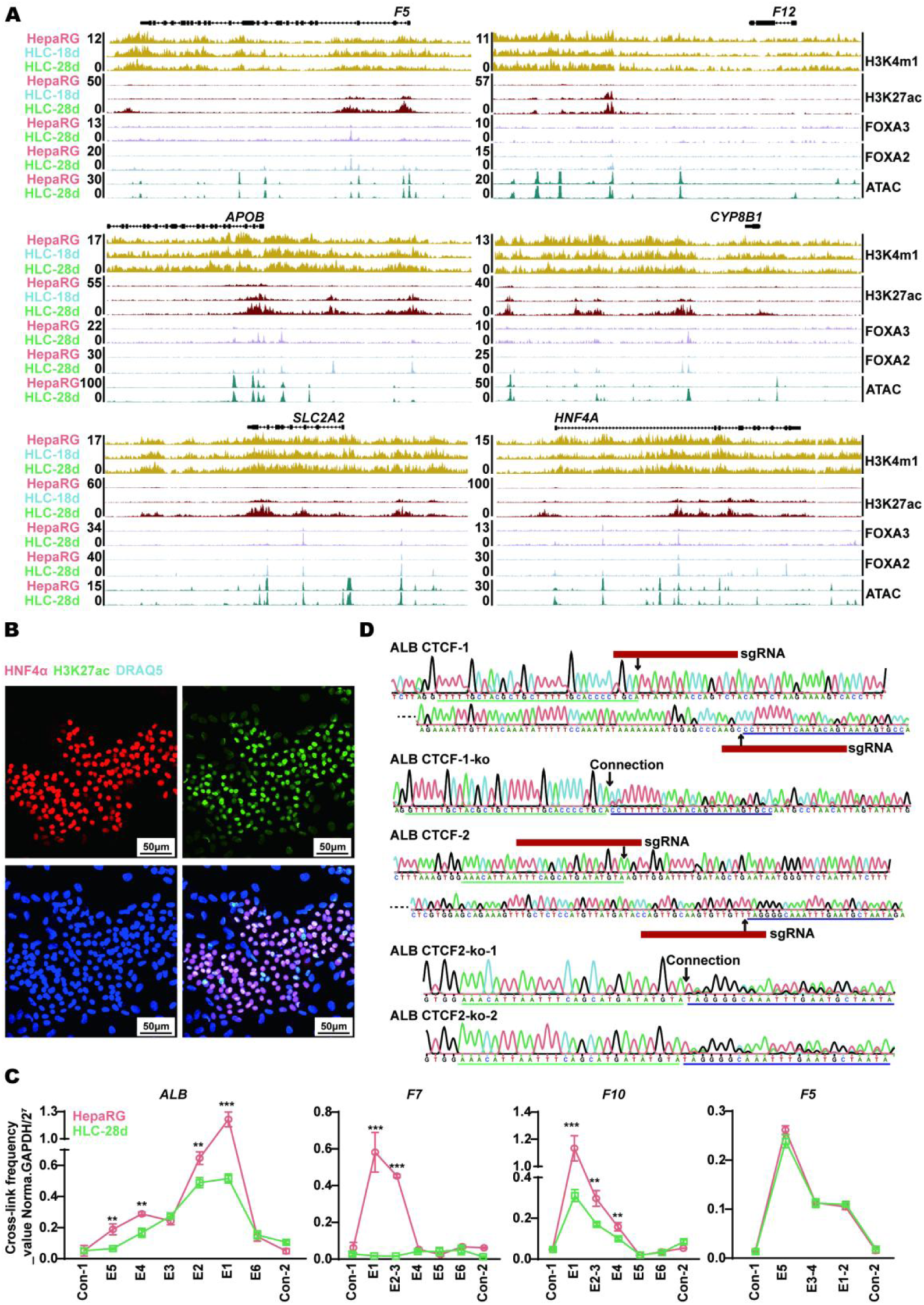
Alterations in epigenetic and chromatin architecture during LPC-to-hepatocyte differentiation. (A) Genome browser tracks based on ChIP and ATAC-seq analyses show transcription factor binding and chromatin modifications at indicated liver function gene loci in HepaRG cells, hepatocyte-like cells at day 18 and day 28. (B) Immunofluorescence staining for HNF4 α (red), H3K27ac (green), and DRAQ5 (blue) was performed and displayed in differentiated HepaRG cells. (C) 3C-qPCR was used to analyze chromatin interaction in liver function genes (ALB, *F5*, *F7*, and *F10*) in undifferentiation HepaRG cells and hepatocyte-like cells. Cross-linking frequency between promoter and enhancer regions of these genes was measured and plotted. Data are shown as mean ± SEM. Statistical significance was determined using appropriate statistical tests: *p < 0.05, **p < 0.01, ***p < 0.001. (D) Sanger sequencing traces confirm CRISPR/Cas9-mediated deletion of CTCF sites in the super-enhancers of the *ALB* gene in undifferentiation HepaRG cells. Following CRISPR editing using specific sgRNAs, targeted deletion of two CTCF binding sites (CTCF-1 and CTCF-2) occurred. Positive clones showed loss of the expected CTCF sequence and altered chromatin connectivity.

**Figure S6.**
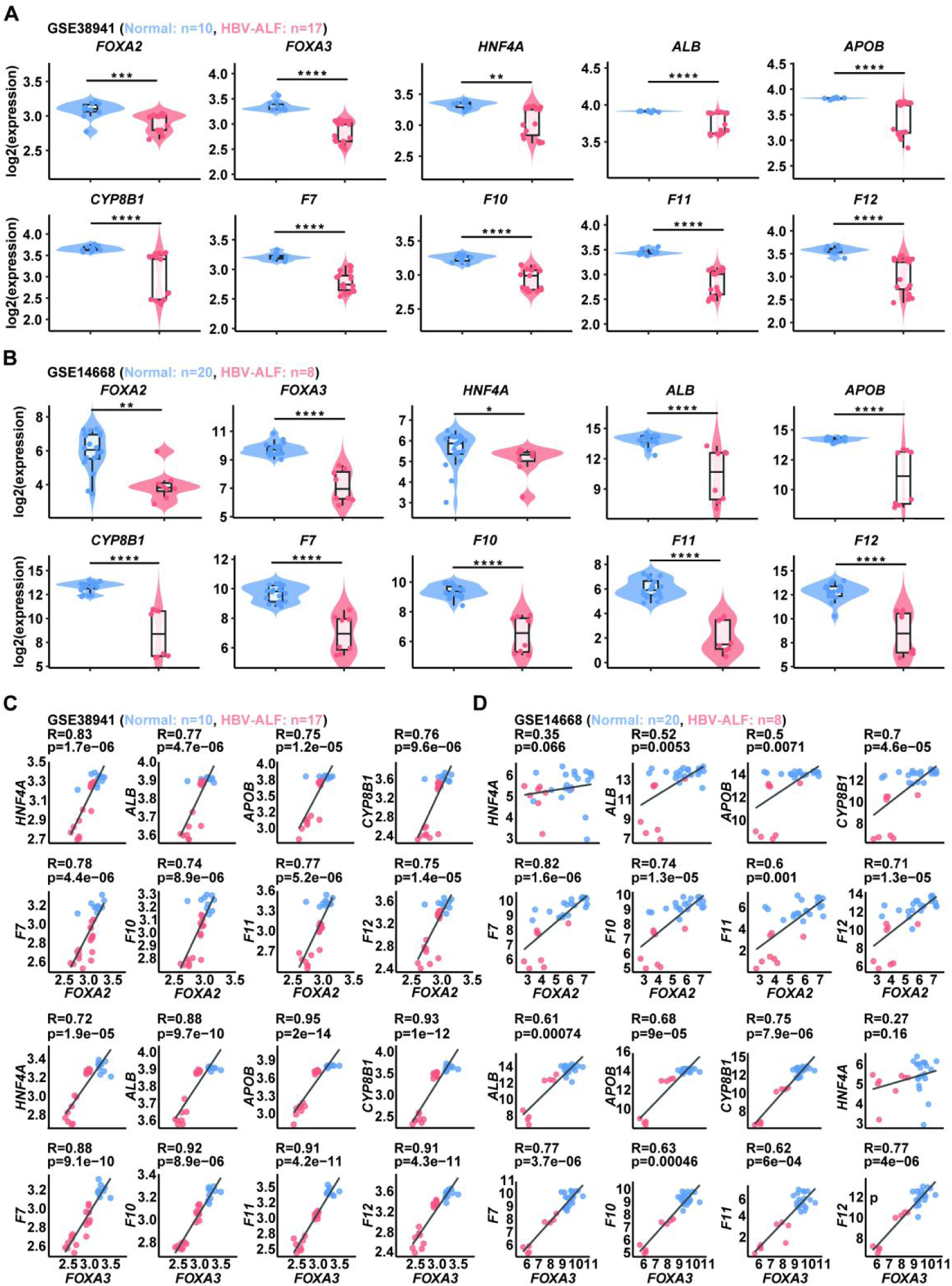
FOXA2 and FOXA3 expression is positively correlated with liver function gene expression in HBV-associated acute liver failure. (**A-B**) Expression profiling by array demonstrates gene expression levels (log2) of liver transcription factors (*FOXA2*, *FOXA3*, *HNF4A*) and liver function genes (*ALB*, *CYP8B1*, *APOB*, *F7*, *F10*, *F11*, *F12*) in healthy control livers (blue) and HBV-ALF patient livers (red). P values were calculated using a two-sided Student’s t-test: *P < 0.05, **P < 0.01, ***P < 0.001, ***P < 0.0001. (**C-D**) Pearson correlation analysis was conducted to assess the relationship between *FOXA2* (top) or *FOXA3* (bottom) and liver function genes with super-enhancers (*ALB*, *APOB*, *CYP8B1*, *F7*, *F10*, *F11*, *F12*, and *HNF4A*) in healthy control livers (blue dots) and HBV-ALF patient livers (red dots). Pearson correlation coefficients (R) and p-values are indicated in each panel.

## References

1 Sheila Sherlock, J. D. Diseases of the Liver and Biliary System. Eleventh edition. (2002).

2 Hall, J. E. Guyton and Hall Textbook of Medical Physiology Twelfth Edition. (2011).

3 David Zakim, T. D. B. Hepatology: a Textbook of Liver Disease, 4th edition. (2003).

4 Hnisz, D. et al. Super-enhancers in the control of cell identity and disease. Cell 155, 934–947, doi:10.1016/j.cell.2013.09.053 (2013).

5 Joo, M. S., Koo, J. H., Kim, T. H., Kim, Y. S. & Kim, S. G. LRH1-driven transcription factor circuitry for hepatocyte identity: Super-enhancer cistromic analysis. EBioMedicine 40, 488–503, doi:10.1016/j.ebiom.2018.12.056 (2019).

6 Lu, X. et al. Super-enhancers in hepatocellular carcinoma: regulatory mechanism and therapeutic targets. Cancer Cell Int 25, 7, doi:10.1186/s12935-024-03599-5 (2025).

7 Weng, H. L. et al. Two sides of one coin: massive hepatic necrosis and progenitor cell-mediated regeneration in acute liver failure. Front Physiol 6, 178, doi:10.3389/fphys.2015.00178 (2015).

8 O’Grady, J. G., Schalm, S. W. & Williams, R. Acute liver failure: redefining the syndromes. Lancet 342, 273–275, doi:10.1016/0140-6736(93)91818-7 (1993).

9 Lefkowitch, J. H. The Pathology of Acute Liver Failure. Adv Anat Pathol 23, 144–158, doi:10.1097/PAP.0000000000000112 (2016).

10 Michalopoulos, G. K. & Khan, Z. Liver Stem Cells: Experimental Findings and Implications for Human Liver Disease. Gastroenterology 149, 876–882, doi:10.1053/j.gastro.2015.08.004 (2015).

11 Roskams, T. et al. Hepatic OV-6 expression in human liver disease and rat experiments: evidence for hepatic progenitor cells in man. J Hepatol 29, 455–463, doi:10.1016/s0168-8278(98)80065-2 (1998).

12 Roskams, T. A. et al. Nomenclature of the finer branches of the biliary tree: canals, ductules, and ductular reactions in human livers. Hepatology 39, 1739–1745, doi:10.1002/hep.20130 (2004).

13 Theise, N. D. et al. The canals of Hering and hepatic stem cells in humans. Hepatology 30, 1425–1433, doi:10.1002/hep.510300614 (1999).

14 Whyte, W. A. et al. Master transcription factors and mediator establish super-enhancers at key cell identity genes. Cell 153, 307–319, doi:10.1016/j.cell.2013.03.035 (2013).

15 Weintraub, A. S. et al. YY1 Is a Structural Regulator of Enhancer-Promoter Loops. Cell 171, 1573–1588 e1528, doi:10.1016/j.cell.2017.11.008 (2017).

16 Hou, Y. et al. Integrative characterization of G-Quadruplexes in the three-dimensional chromatin structure. Epigenetics 14, 894–911, doi:10.1080/15592294.2019.1621140 (2019).

17 Dowen, J. M. et al. Control of cell identity genes occurs in insulated neighborhoods in mammalian chromosomes. Cell 159, 374–387, doi:10.1016/j.cell.2014.09.030 (2014).

18 Dixon, J. R., Gorkin, D. U. & Ren, B. Chromatin Domains: The Unit of Chromosome Organization. Mol Cell 62, 668–680, doi:10.1016/j.molcel.2016.05.018 (2016).

19 Lucke, B. & Mallory, T. The fulminant form of epidemic hepatitis. Am J Pathol 22, 867–947 (1946).

20 Lin, T., Feng, R., Liebe, R. & Weng, H. L. Liver Progenitor Cells in Massive Hepatic Necrosis-How Can a Patient Survive Acute Liver Failure? Biomolecules 12, doi:10.3390/biom12010066 (2022).

21 Cerec, V. et al. Transdifferentiation of hepatocyte-like cells from the human hepatoma HepaRG cell line through bipotent progenitor. Hepatology 45, 957–967, doi:10.1002/hep.21536 (2007).

22 Lin, T. et al. Follistatin-controlled activin-HNF4alpha-coagulation factor axis in liver progenitor cells determines outcome of acute liver failure. Hepatology 75, 322–337, doi:10.1002/hep.32119 (2022).

23 Blobel, G. A., Higgs, D. R., Mitchell, J. A., Notani, D. & Young, R. A. Testing the super-enhancer concept. Nat Rev Genet 22, 749–755, doi:10.1038/s41576-021-00398-w (2021).

24 Feng, R. et al. A hierarchical regulatory network ensures stable albumin transcription under various pathophysiological conditions. Hepatology 76, 1673–1689, doi:10.1002/hep.32414 (2022).

25 Li, J., Ning, G. & Duncan, S. A. Mammalian hepatocyte differentiation requires the transcription factor HNF-4alpha. Genes Dev 14, 464–474 (2000).

26 Lemaigre, F. P. Mechanisms of liver development: concepts for understanding liver disorders and design of novel therapies. Gastroenterology 137, 62–79, doi:10.1053/j.gastro.2009.03.035 (2009).

27 Kyrmizi, I. et al. Plasticity and expanding complexity of the hepatic transcription factor network during liver development. Genes Dev 20, 2293–2305, doi:10.1101/gad.390906 (2006).

28 Iwafuchi-Doi, M. & Zaret, K. S. Pioneer transcription factors in cell reprogramming. Genes Dev 28, 2679–2692, doi:10.1101/gad.253443.114 (2014).

29 Brown, J. D. et al. NF-kappaB directs dynamic super enhancer formation in inflammation and atherogenesis. Mol Cell 56, 219–231, doi:10.1016/j.molcel.2014.08.024 (2014).

30 Godoy, P. et al. Extracellular matrix modulates sensitivity of hepatocytes to fibroblastoid dedifferentiation and transforming growth factor beta-induced apoptosis. Hepatology 49, 2031–2043, doi:10.1002/hep.22880 (2009).

31 Hagege, H. et al. Quantitative analysis of chromosome conformation capture assays (3C-qPCR). Nat Protoc 2, 1722–1733, doi:10.1038/nprot.2007.243 (2007).

32 Thakore, P. I. et al. Highly specific epigenome editing by CRISPR-Cas9 repressors for silencing of distal regulatory elements. Nat Methods 12, 1143–1149, doi:10.1038/nmeth.3630 (2015).

33 Sanjana, N. E., Shalem, O. & Zhang, F. Improved vectors and genome-wide libraries for CRISPR screening. Nat Methods 11, 783–784, doi:10.1038/nmeth.3047 (2014).

